# Multiomic and quantitative label-free microscopy-based analysis of *ex vivo* culture and TGFbeta1 stimulation of human precision-cut lung slices

**DOI:** 10.1101/2019.12.13.875963

**Authors:** Muzamil Majid Khan, Daniel Poeckel, Aliaksandr Halavatyi, Frank Stein, Johanna Vappiani, Daniel C. Sevin, Christian Tischer, Nico Zinn, Jess D Eley, Natasja Stæhr Gudmann, Thomas Muley, Hauke Winter, Andrew J Fisher, Carmel B. Nanthakumar, Giovanna Bergamini, Rainer Pepperkok

## Abstract

Fibrosis can affect any organ resulting in the loss of tissue architecture and function with often life-threatening consequences. Pathologically, fibrosis is characterised by expansion of connective tissue due to excessive deposition of extracellular matrix proteins (ECM), including the fibrillar forms of collagen. A significant hurdle for discovering cures for fibrosis is the lack of suitable models and techniques to quantify mature collagen deposition in tissues. Here we have extensively characterized an ex-vivo cultured human lung derived, precision-cut lung slices model (hPCLS) using live fluorescence light microscopy as well as mass spectrometry-based techniques to obtain a proteomic and metabolomic fingerprint. Using an integrated approach of multiple readouts such as quantitative label-free Second Harmonic Generation (SHG) imaging to measure fibrillar collagen in the extracellular matrix and ELISA-based methods to measure soluble ECM biomarkers, we investigated TGFbeta1-mediated pro-fibrotic signalling in hPCLS. We demonstrate that hPCLS are viable and metabolically active with mesenchymal, epithelial, endothelial, and immune cells surviving for at least two weeks in ex vivo culture. Analysis of hPCLS-conditioned supernatants showed strong induction of ECM synthesis proteins P1NP and fibronectin upon TGFb stimulation. Importantly, this effect translated into an increased deposition of fibrillar collagen in ECM of cultured hPCLS as measured by a novel quantitative SHG-based imaging method only following addition of a metalloproteinase inhibitor (GM6001). Together the data show that an integrated approach of measuring soluble pro-fibrotic markers and quantitative SHG-based analysis of fibrillar collagen is a valuable tool for studying pro-fibrotic signalling and testing anti-fibrotic agents.

## Introduction

Excessive deposition of extracellular matrix proteins is a hallmark of fibrosis. This leads to alteration of tissue architecture, subsequently loss of its function and ultimately to end-stage organ failure^1^. In the developed world, 45% of all deaths are attributed to conditions related to an excess of ECM deposition^2^. While fibrosis contributes to exacerbated pathology in cancer, myocardial infarctions and ageing, it is also a primary cause of mortality in fibrotic disease of kidneys, liver and lung in particular^3^. Chronic interstitial lung disease (ILDs) is the most common form of pulmonary fibrosis (*European Respiratory Society*). ILDs have been subdivided into over 300 subtypes among which idiopathic pulmonary fibrosis (IPF) is the most prevalent (*European Respiratory Society*). Upon damage of epithelial and endothelial cells in various organs, an inflammatory response is launched which triggers blood clot formation and ECM repair. Part of this repair mechanism is the release of cytokines such as TGFß1 that initiate the activation of macrophages and fibroblasts^3^. Activated fibroblasts express alpha-SMA, which leads to their differentiation in myofibroblasts^3^. Persistent chronic inflammation triggers unchecked proliferation of myofibroblasts, increased epithelial to mesenchymal cell type transition ^3^. Mesenchymal population of cells have an enhanced ability to produce ECM components and a combination of these events result in a derailed tissue repair process leading to tissue scarring^3^. TGFß1 is considered to be the master regulator of signalling pathways (pro-fibrotic) that leads to an abnormally excessive deposition of extracellular matrix^4^. Induction of pro-fibrotic signalling also shifts the delicate balance between synthesis and breakdown of collagens and other ECM^5^ proteins in favour of excessive deposition and pathological stiffening of ECM with loss of tissue compliance. This excessive deposition and stiffness of ECM causes a physical abnormality that leads to atrophying of alveolar tissue^3^ and subsequent suffocation. In IPF patients the fibrotic foci are characterized by abnormally high content of fibrillar collagen (type I, III and V)^3,6^. At the ECM level, presence of these fibrillar collagens are the physical cause of eventual mortality^7^. Despite our understanding of the molecular basis underlying the fibrotic process, to date only two licensed drugs; pirfenidone and nintedanib^8^ are approved for the treatment of IPF. These medicines have very low efficacy and hence patient high withdrawal rates^9^. The lack of more effective anti-fibrotic therapies may be attributed in part (a) to the absence of adequate model systems that can mimic the pathophysiology of the fibrotic process as related to human physiology^10^, (b) to indirect functional readouts of biomarkers used for quantifying disease progression. Currently, using ELISA based measurements of cleavage products of collagen propeptides such as type I or III and VI, collagen can be quantified as they are cleaved of during formation. These cleavage products reveal information on fibrosis disease progression^11^. While these biomarkers are crucial their correlation/ dynamics to excess deposition is still to be established. Microscopic detection of fibrillar collagen is also difficult due to lack of specific antibodies. Therefore, we have sought to establish a quantitative label-free second harmonic generation imaging workflow for fibrillar collagen in hPCLS. Murine experimental models of fibrosis are frequently used to study lung, liver and kidney fibrosis. Typically, pulmonary fibrosis is induced either using genetic manipulation (MUC5b overexpression, a genetic risk factor identified in large cohorts of IPF patients)^12^ or chemical induction (bleomycin sulphate, amiodarone, carbon tetrachloride)^13^. While these model systems help to understand acute signalling pathways preceding tissue damage and, in some instances, chemicals such as bleomycin sulphate can be used as causative agents^14^. Furthermore, *in vivo* models lack histological features of human disease and can spontaneously resolve, therefore failing to recapitulate the irreversible end-stage disease. These model systems have highlighted an inflammatory component present during the development of fibrotic disease, yet anti-inflammatory drugs have failed in clinical development^15,16^. Based on these findings, therapies involving corticosteroids (prednisone) with or without immunosuppressive drugs (azathioprine and cyclophosphamide) were used to treat IPF. In fact, some drugs have had adverse effects that worsen patient condition^15,16^.

Human precision-cut lung slices (hPCLS), 3-dimensional, uniformly cut slices of lung tissue that may be maintained in *ex vivo* culture have emerged as a promising potential model system. PCLS (rat lung and human) came into existence in the early 1990s^17,18^ but have only recently been used as a model for evaluating human disease pathophysiology^19^. Several groups have used this model system to recapitulate pro-fibrotic signalling^10,20,21^although the mechanisms at play and the validation of the molecular changes that hPCLS undergo in *ex vivo* culture are still elusive. These studies do show an induction in fibrotic signalling but a functional readout of enhanced ECM deposition is lacking. Furthermore, none of these studies have characterized the presence of fibrosis essential cell types such as epithelial, mesenchymal and immune cell types^3^. Myofibroblasts are the key cellular source of collagen in fibrosis but these cells are derived from tissue-resident mesenchymal, epithelial and endothelial cells^3^.

Also, it is assumed that in hPCLS cell/cell and cell/matrix interactions are well preserved, however, no study has been conducted to evaluate such claims. Therefore, in this study using mass spectrometry and light microscopy-based approaches we first sought to investigate and characterize the changes of protein expression in the four major cell types present in lung tissue when hPCLS are *ex vivo* cultured. Second, to investigate the pro-fibrotic signalling induced by TGFß1 stimulation, we established a new, quantitative, label-free second harmonic generation imaging method that can measure deposited fibrillar collagen in the extracellular matrix upon pro-fibrotic stimulation. This model represents a human translational model system that could be used for testing antifibrotic agents.

## Results

### Proteomic dissection of molecular changes in hPCLS over time in *ex-vivo* culture conditions

To use hPCLSs for mimicking a particular disease involving certain cell types and signalling pathways, it is imperative to characterize this model system in given *ex vivo* culture conditions. We used mass spectrometry-based proteomics and untargeted metabolomics to monitor hPCLS proteome and metabolome over time in *ex vivo* culture. Here, *ex vivo* cultured hPCLS in DMEM were harvested on different days (day 1, day 4, day 7, day 10 and day 13) and snap-frozen in liquid nitrogen. Subsequently, lysates were prepared for mass spectrometry-based proteomic and metabolomic analysis. For proteomic analysis, 2 % SDS soluble fraction of hPCLS (4 donors) were subjected to LC/MS analysis. Approximately 5500 proteins were detected across 4 donors (2 × 2hPCLS/donor at each time point) and of those, 4288 common proteins were subjected to analysis (excel sheet-SuplementaryProtTimeCourse). Differential analysis (Figure 1 A) of log2 fold change in protein levels with respect to the previous time point of harvest was plotted. The analysis shows (blue and red points) that hPCLS undergo moderate molecular changes compared to day 1 till day 13 (in total on day 13, 17%, 728 out of 4288 proteins changed significantly). Of the 17% changes, 10% of changes take place from day 1 to day 7 and the remaining 7% from day 7 to day 13. No new proteins fit the significance criterion upon comparing day 10, day 13 to day 7 and 10 respectively (Figure1A). On day 13, 343 proteins out of 728 regulated genes were downregulated and 385 were upregulated.

**Figure 01:**
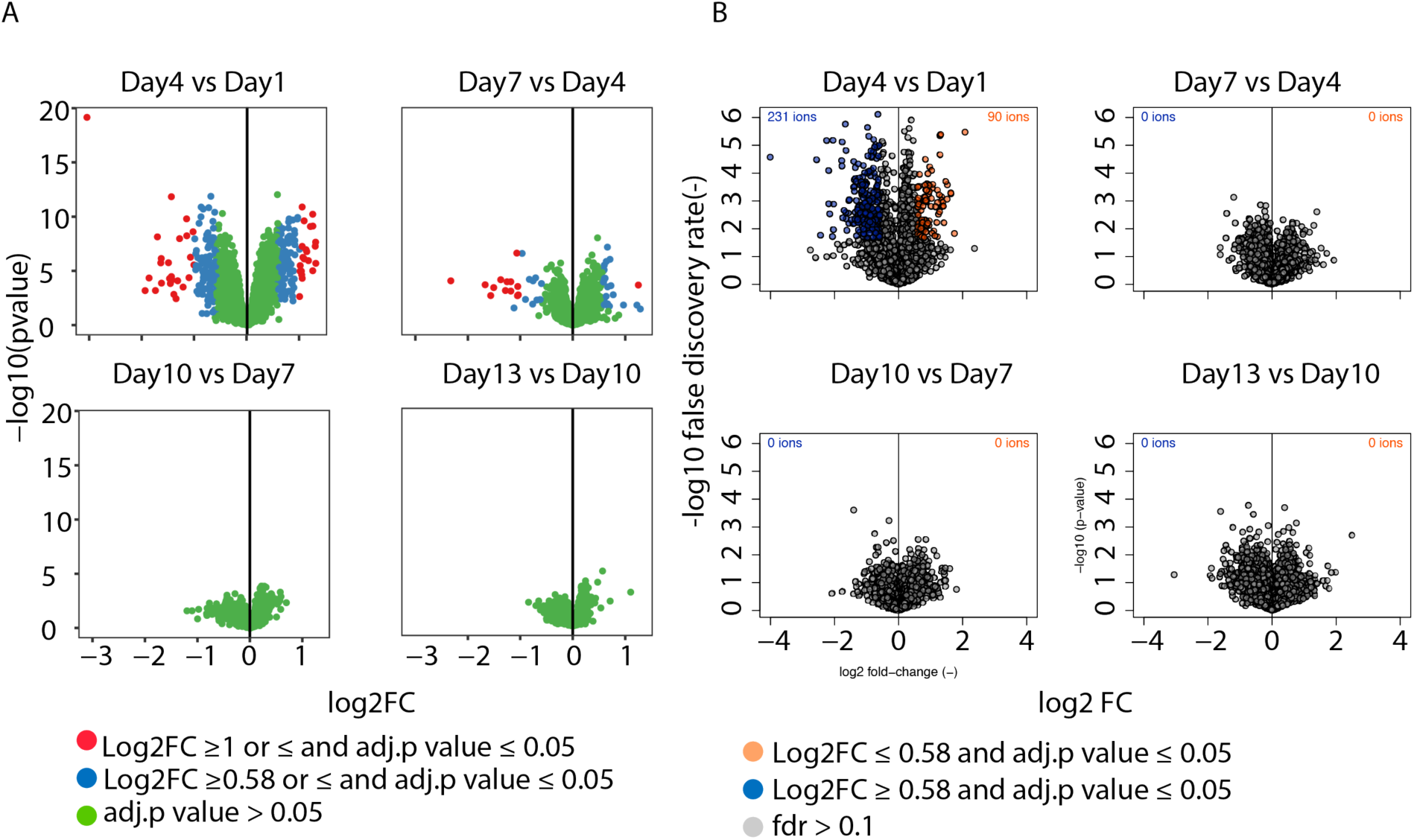
Multiomic analysis of molecular changes in ex vivo cultured hPCLS: (A) Volcano plot analysis of the proteome of hPCLS cultured ex vivo over time. Log2FC of the respective proteome on a given day of culture was normalized to the previous time point. Analysis highlights that hPCLS undergo moderate changes in first few days of culture and remain stable over time. (B) Untargeted metabolomic analysis of hPCLS was performed. Volcano plot analysis of log2FC changes in ions was performed similarly as above. The analysis confirms that hPCLS undergo very moderate metabolic changes similarly as the proteome changes.

Cytoscape based network analysis (Supplementary Figure 1B) of significantly regulated proteins on day 13 vs day 1 showed primarily an upregulation of pathways involving proteins related to degrading ECM whereas only few pertaining to the formation of ECM. A significant upregulation of inflammatory signalling pathways were observed, such as neutrophil degranulation and innate immune system.

We further analysed the proteomics data to investigate the dynamics of different cell types underlying the observed changes in protein levels. Human lung tissue is mainly composed of Immune, mesenchymal, endothelial and epithelial cells. **L**ung **G**ene **E**xpression **A**nalysis (LGEA) web portal has established a database of gene expression of each cell type (also age-dependent, from neonatal to adult stages) in both mouse and human lungs^22,23^. In this database, the transcriptome of flow-sorted human lung cells have been designated as CD45+ immune cells, CD45-/PECAM-/VECadherin-/EpCAM+ mixed epithelial cells, CD45-/PECAM+/VECadherin+ mixed endothelial cells and CD45-/PECAM-/VECadherin-/EpCAM-mixed mesenchymal cells. We compared (excel sheet-SuplementaryCellTypeComp-TimeCourse) significantly regulated (log2FC ≥0.5 and ≤-0.5 and fdr p-value ≤ 0.05) proteins on day 13 vs day 1 to the list of exclusive proteins attributed to different cell classes (in LGEA^23^ analysis). Figure Suppl. 01C shows that different cell type markers are present on day 13 in our culture system, and no exclusive upregulation nor any downregulation of a particular cell type could be identified by this method over the time here analyzed. Of the 728 significantly regulated proteins (day 13 vs day 1), we found proteins exclusively expressed in specific cell types, 61 in mesenchymal, 33 in immune, 25 in epithelial and 32 in endothelial cells. Among the significantly regulated mesenchymal proteins, pro-fibrotic collagens^7^ such COL5A1 (Log2FC −0.7) and COL3A1 (Log2FC −0.9) were significantly downregulated. Given collagen class of proteins is highly relevant for fibrosis progression^7^, this observation prompted us to look into the pattern of regulation of all the collagens detected. Data shows that under current culture conditions, the majority of the detected collagens (TableS1) undergo downregulation over time including COL1, COL4 and COL6. Furthermore, there is a significant upregulation of MMP2 (Log2FC 2.1) and MMP14 (Log2FC 2.27), metalloproteinases involved in degrading fibrillar collagens^24^ such as COLI, COLII, COLIII and COLV. In general, the metalloproteinases detected over the course of ex vivo culture show an upregulated expression (Table S1).

### Metabolomic and live imaging analysis hPCLS stability over time in *ex-vivo* culture conditions

To investigate whether hPCLS are metabolically active ex vivo, samples were harvested as for the proteomic analysis, and in addition solubilized and mechanically homogenized to extracts metabolites. These were subjected to LC/MS-based untargeted metabolomic analysis ^25^. A total of 3523 metabolites were detected (excel sheet-Suplementarymetabolome) reproducibly in > 90% of the samples (3 donors, 3 technical replicates/ donor, 1 technical replicate=2 hPCLS). Contrary to proteomics data that showed a linear increase in significantly regulated proteins (each day compared day 1), metabolome had ca. 330 metabolites differentially regulated on day 04, 07, 10 and 13 (compared day 1). The differential analysis of metabolomics data (Figure. 1B) revealed that the hPCLS metabolome undergoes changes between day 1 to day 04 of *ex vivo* culture but is subsequently stable with no statistically significant changes from day 04 to day 13 (Figure1B). Similar to the proteomic data, metabolomic data also show that the changes occurred initially between day 1 to day 04 increase in magnitude till day 13 (supplementary Figure 01E). We also performed a pathway analysis of the significantly regulated annotated metabolites (day 1 vs day 13), but no enrichment of specific pathways was observed. We analysed changes in the response of selected metabolites sensitive to changes in tissue health. Here we observed hPCLS are metabolically stable in *ex vivo* culture (Supplementary Figure01 F). Of note, Adenosine triphosphate levels of the hPCLS remained unchanged and L-Lactic acid levels significantly built up over time in culture, implicating a metabolically active 3D tissue system. Also, L-Glutamine was increasingly being metabolized by hPCLS without undergoing any oxidative stress (as depicted by stable levels of the redox-sensitive metabolites cysteine-glutathione disulfide and glutathione).

To further assess cell viability in hPCLS ex vivo culture we also performed live confocal microscopy for live/ dead cell analysis (CalceinAM/ Ethidium homodimer kit). Subsequent image analysis showed that there is no significant change in live-cell signal over time in the *ex vivo* culture from day 1 to day 13 (Supplementary Figure01A) while dead cell number considerably decreased from day 1 to day 04. These results are consistent with other studies^10,20^ that have assessed the viability of hPCLS *ex vivo* culture albeit for 5-7 days.

Taken together these data show that in this hPCLS *ex-vivo* culture system all the relevant cell types (mesenchymal, epithelial, endothelial, immune) and signalling pathways whose interplay is important for studying pro-fibrotic signalling, are present. The system shows the presence of complex healthy wound healing response upon exposure to physical injury (tissue slicing), and of inherent metalloproteinase signalling, extracellular matrix organization and smooth muscle responses (Supplementary Figure 01B).

### Induction of pro-fibrotic signalling upon TGFß1 stimulation in *ex-vivo* hPCLS culture

TGFß1 is a cytokine regarded as master regulator of fibrotic signalling^26^. To induce pro-fibrotic signalling in our culture system we used TGFß1 (10ng/ml) as reported previously by other groups both in murine model systems^27^ and hPCLS^10^, however in our case for longer time and repetitively. Here hPCLS were replenished with fresh medium containing human recombinant TGFß1 after every 72 hours for 13 days in ex vivo culture. Supernatants were collected on day 4, day 7 and day 13. These supernatants were subjected to proprietary competitive ELISA for detecting selected neoepitope^28^ pro-fibrotic markers such as Procollagen 1 N-terminal propeptide (PRO-C1)^29^, the C-terminal part of Fibronectin (FBN-C), Collagen type III degradation marker (C3M). Consistent with other studies^10,30^, TGFß1 stimulation increased the level of PRO-C1 and FBN-C detected in the media supernatants indicating induction of pro-fibrotic signalling, while C3M showed no changes. To better characterize how the hPCLS model system would respond to TGFß1 stimulation, we analysed TGFß1 effects at the proteomic level (4 donors). Here, hPCLS (vehicle-treated and TGFß1 treated) were harvested on day 13 of *ex vivo* culture. The hPCLS were subjected to 2% SDS solubilization and mechanical disruption. Subsequently, lysates were subjected to LC/MS. A total of 6600 proteins per replicate per donor were detected, of these only 3998 common proteins found (excel sheet SuplementaryProtTGF) in 8 replicates (4 donors) were subjected to downstream data analysis. On day 13 hPCLS treated with TGFß1 showed 109 proteins were significantly (Log2FC between 0.58 and −0.58, fdr p-value of 0.05) regulated (out of 3998, Figure supplement 2A). A curated data analysis of these proteins showed upregulation of prominent pro-fibrotic markers such as COL1A2, COL3A1, COL5A1, FBN-C and many others (Figure 2B). Consistent with other studies many TGFß1 regulated pro-fibrotic markers such as THBS1^31^, THBS2^32^, VCAN and TNC^33^, FKBP10^34^ were upregulated as well. Furthermore, significantly regulated proteins were subjected to pathway analysis using Cytoscape (Supplementary Figure2B). This analysis shows induction of pro-fibrotic pathways such as embryonic morphogenesis, ECM organization, interleukin signalling and downregulation of pro-epithelial injury signalling pathways such as surfactant metabolism^35–38^. Interestingly, proteomic analysis of peripheral blood from IPF patients has been shown to have downregulation of cell proliferation^39^pathways, similar to the proteome of TGFß1 treated hPCLS analysed in our study.

**Figure 02:**
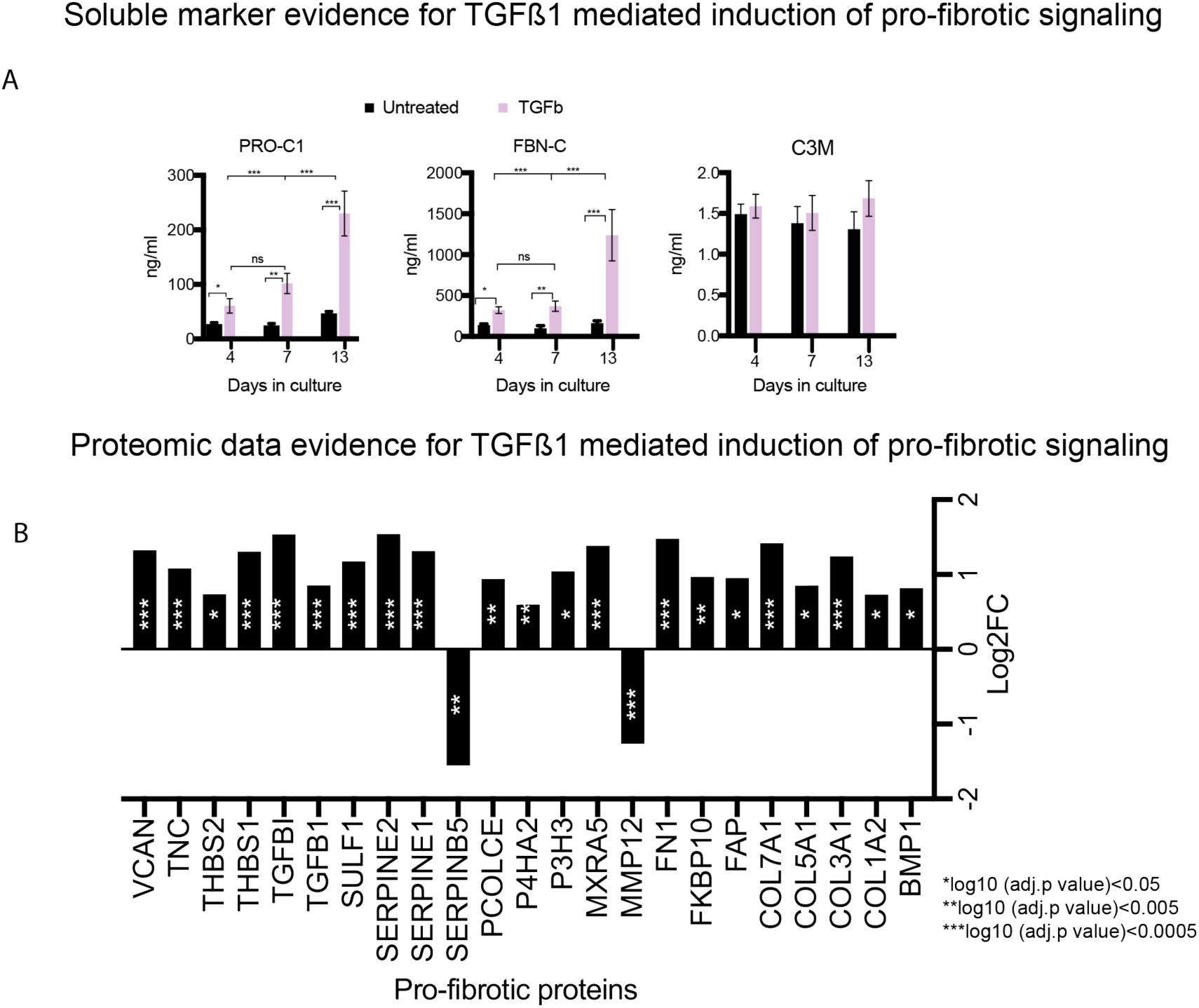
Assessment of pro-fibrotic signalling induction upon TGFß1 stimulation of ex vivo cultured hPCLS: (A) ELISA analysis of media supernatants of vehicle and TGFß1 treated hPCLS. Here PRO-C1, Fibronectin-C peptide and C3M soluble pro-fibrotic markers were analysed. Bar graphs represent an average of 4 media supernatants (each from a hPCLS). (B) Log2FC of pro-fibrotic curated protein markers in mass spec data upon TGFß1 stimulation on day 13 (with respect to day 1) of hPCLS culture. Results show clear upregulation of majority of these markers. ELISA error bars SEM, *p<0.05, **p< 0.01***p<0.001, ANOVA with Turkey’s multiple comparisons test. Proteomics datap-values represent false discovery rate.

Furthermore, we curated the significantly regulated proteins on day 13 in TGFß1 treated hPCLS (vs day 13, unstimulated hPCLS) to those of different cell type markers listed in LGEA web portal^22,23^. Strikingly, data (supplementary figure 2C) shows that the majority of the significantly downregulated proteins (ca. 88%, 39 out of 44) belong to Epithelial class of cells while those upregulated belong to mesenchymal cell class (ca. 99%, 64 out of 65). These data support that this TGFß1 stimulation induces an epithelial to mesenchymal transition based pro-fibrotic signalling system^4^ in our culture condition of hPCLS. Both readouts of collagen synthesis, ELISA for the supernatant and SDS soluble fraction for the intracellular, showed high upregulation (Figure 2A-B). While it is important to measure regulation of various markers of a pro-fibrotic process to validate induction of pro-fibrotic signalling, to better assess the functional consequences it is necessary to confirm that this signalling translates into higher extracellular deposition of fibrillar collagen. Several recent studies^10,20,21^ have shown induction of pro-fibrotic signalling in *ex-vivo* cultured hPCLS using TGFß1 stimulation^10^ or a cocktail of pro-fibrotic factors^20^ but a validation of consistent and specific ECM deposition (fibrillar collagen in particular) is still lacking. Next we investigated whether the observed upregulation was translating into deposition of fibrillar collagens in the extracellular matrix.

### Quantitative label-free Second Harmonic Generation (SHG) imaging analysis of fibrillar collagen in ECM upon TGFß1 stimulation

Here, we employed label-free SHG imaging to quantitatively measure deposition of fibrillar collagen in our ex vivo cultured hPCLS upon TGFß1 stimulation. When biological tissue is illuminated with 2 photon laser, individual noncentrosymmetric molecules generate a second harmonic signal with double the frequency and half the wavelength length of the input laser^40^. In human tissues acto-myosin complex of skeletal muscles (cytosolic)^41^, large microtubules^42^ (cytosolic) and fibrillar collagen^40^ (extracellular matrix) have the noncentrosymmetry required to generate SHG signal. Given, large fibrillar collagen is only present in ECM, it makes SHG imaging a highly specific and a suitable technique for quantitative measurements of matrix deposition. Even though SHG imaging of tissue has been employed in several studies ^43–45^ and also in murine models of pulmonary fibrosis^12^, a systematic quantitative workflow in particular for human lung tissue samples is still lacking. First, we compared healthy hPCLS (derived from tumour-free lung tissue) with those derived from end-stage interstitial lung disease [(Idiopathic pulmonary fibrosis (IPF) and Nonspecific interstitial pneumonia (NSIP)] patients.

patients using SHG imaging. 250µm thick and 8mm in diameter hPCLS from day 0 were chemically fixed. The entire hPCLS were imaged (Figure 3A-B). Subsequently, the acquired 3D stacks were analysed using a semi-automated image analysis pipeline (Jython script-image analysis) in FIJI. As expected, the analysis showed significantly more deposited fibrillar collagen in ILD hPCLS. We observed the SHG signal originating from airways or blood vessels (Figure 3B*) was similar in intensity between healthy and ILD tissues, while interstitial collagen signal (Figure 3B*) showed dramatic increase in ILD compared to healthy condition. Next, to investigate whetherTGFß1 induced pro-fibrotic signalling (Figure 2A-B) is translated into excessive deposition of collagen in lung ECM, vehicle-treated and TGFß1 treated hPCLS were similarly chemically fixed with 4% PFA on day 13 of ex vivo culture. SHG imaging of whole slices was performed. However, high variability was observed from the analysis within untreated or treated hPCLS as well across two conditions. After careful observation, it became evident that this variability originates from fibrillar collagen present in different structures of hPCLS (Supplementary Figure 3). Different hPCLS have variable amounts SHG signal originating from the constituent fibrillar collagen of blood vessel or airways and that of lung interstitium (Supplementary Figure 3A). Therefore, to focus our analysis to interstitium of the hPCLS, we applied a histological approach^46^ of image analysis where the 3D SHG image stacks of hPCLS were divided into equally sized grids or regions of interest. Using a semi-automated FIJI based script, we manually excluded any grid that has SHG signal originating from a blood vessel or an airway was excluded (purple Xs, Figure 3A) from analysis, hence, only interstitium of the hPCLS was analysed. Furthermore, it must be noted for analyzing fibrotic phenotype in interstitial lung diseases it is the parenchymal interstitium^47^ which is affected, hence the need to analyze the interstitium. This approach is further supported by our observation of increased SHG signal of interstitium in IPF vs. healthy hPCLS (Figure 3B-C). This analysis shows that there is a marginal increase in the deposition of fibrillar collagen upon TGFß1 stimulation but not statistically significant (Figure 3E).

**Figure 03:**
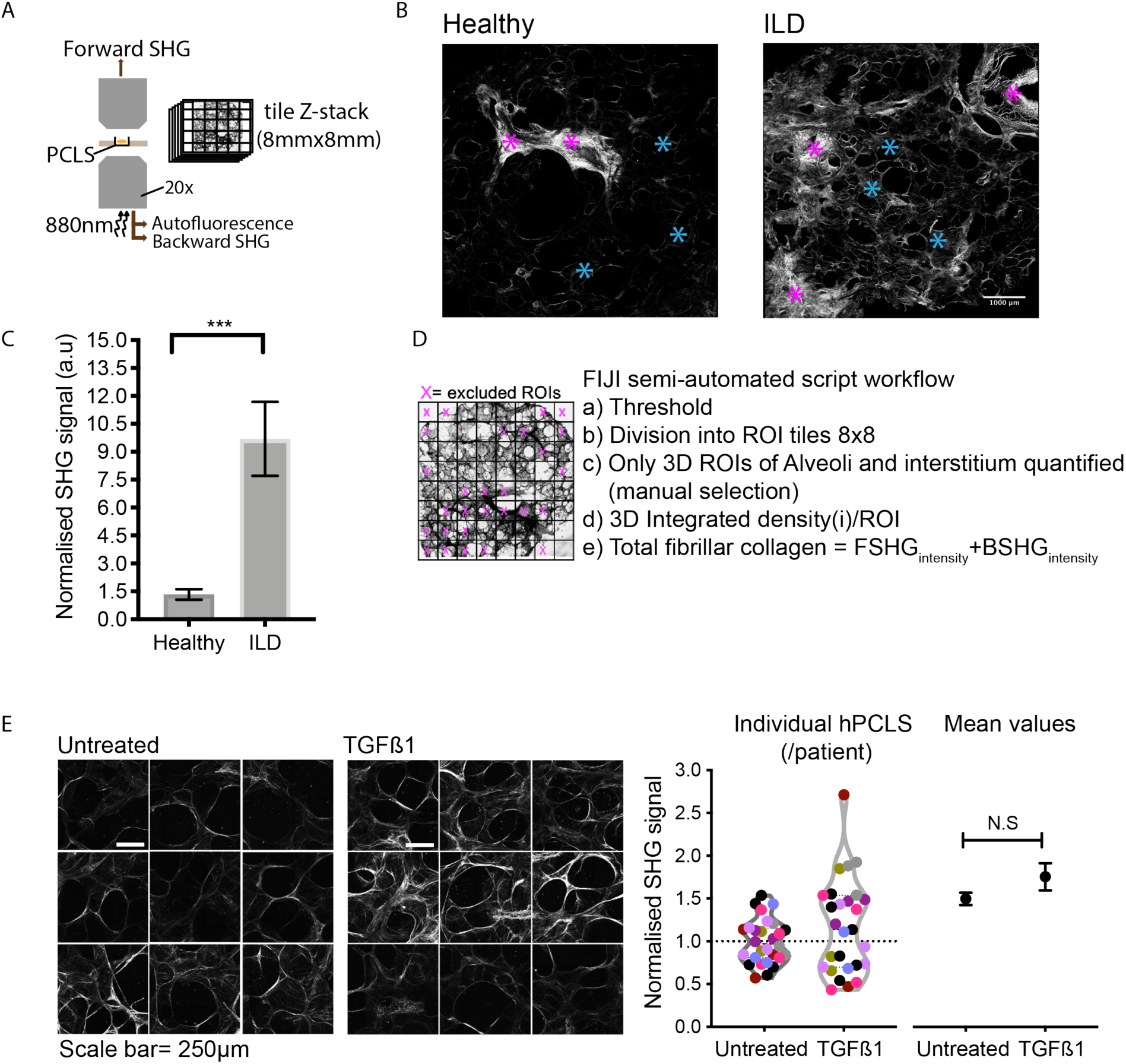
Second Harmonic generation image analysis of fibrillar collagen deposition in ex vivo cultured hPCLS upon TGFß1 stimulation: (A) The scheme represents the SHG microscopy and imaging setup. hPCLS are images in a 3D tiled scan with a 20X air objective. Semi-automated FIJI based script divides the SHG 3D image stack of hPCLS into 8×8 grids (approx. 0.8X0.8mm). Magenta “X” represents the example rois that excluded from the analysis. (B-C) Maximum Z-projection and quantification of SHG z-stacks of healthy and ILD hPCLS (integrated SHG signals normalized to average healthy condition) (D) workflow for quantifying SHG signal originating from lung interstitium (E) Maximum Z-projection representative rois of vehicle-treated and TGFß1 treated hPCLS (same donor). Forward and backward SHG channel intensity is added and total fibrillar collagen is calculated. SHG intensity of TGFß1 rois is normalized to the mean SHG intensity of same donor vehicle-treated hPCLS. Color of the points in the violin plots represent hPCLS/ donor. Mean normalized SHG intensity signal value of hPCLS is shown. Colored dots represent each hPCLS/ donor (8 donor with 3-4 hPCLS/ donor). Two tailed Students t-test was performed for healthy and ILD SHG signal comparison(C), ***p=0.0008. Students t-test between means of vehicle and TGFß1 treated hPCLS was performed. No significance (N.S) was observed using two tailed students t-test.

### Metalloproteinase (MMP) inhibitor together with TGFß1 stimulation significantly enhances deposition of fibrillar collagen in ECM

The proteomics analysis showed that untreated hPCLS over time in *ex vivo* culture show a wound healing response with a fine balance between construction and degradation of ECM (Supplementary Figure 1B). Therefore, even though a pro-fibrotic signal is induced, this induction might not be enough for conferring deposition of fibrillar collagen in the ECM of this hPCLS *ex vivo* model.

Furthermore, studies have shown that the excessive deposition fibrillar collagen in the ECM of fibrotic tissues is due to an imbalance of proteins involved in ECM formation and degradation. A major component of this imbalance is that of abnormal MMP activity in fibrotic tissues^48^. Furthermore, in murine models of PCLS, collagen type I, III and elastin have been shown to have high turnover due to higher MMP activity^49^. Han S. et al.^50^ has shown that pan inhibition of MMP activity using Ilomastat (GM6001) enhances collagen fibril formation in human mesenchymal stem cell *in vitro* culture. Encouraged by these observations, hPCLS were concomitantly treated with Ilomastat and TGFß1 for two weeks. Subsequently, supernatants of hPCLS (Vehicle, TGFß1 and TGFß1+GM6001) were collected and subjected to ELISA measurements for PRO-C1 and FBN-C. Data show that concomitant treatment of hPCLS with TGFß1 and Ilomastat significantly increases the levels of both PRO-C1 and FBN-C in the media supernatants (Figure 4C) in comparison to TGFß1 treatment alone. This observation can be attributed to reduced degradation of these proteins due to MMP inhibition. It is important to note that Ilomastat increased the levels of PRO-C1 approximately 3.5 times while as the effect on FBN-C was moderate. Since, MMP inhibition resulted in an enhanced population of PRO-C1 available for deposition, we again analysed PFA fixed hPCLS on day 13 of *ex vivo* culture (vehicle, TGFß1 and TGFß1+GM6001), The SHG analysis of these hPCLS showed a significantly increased deposition of fibrillar collagen in the ECM (Figure 4A-B), indicating that in the conditions here optimized for culturing hPCLS, ECM deposition can be quantitatively measured by SHG. It should be noted that hPCLS derived from two patients (magenta and yellow points Figure 4B) did not show higher deposition upon concomitant treatment of TGFß1 and GM6001.

**Figure 04:**
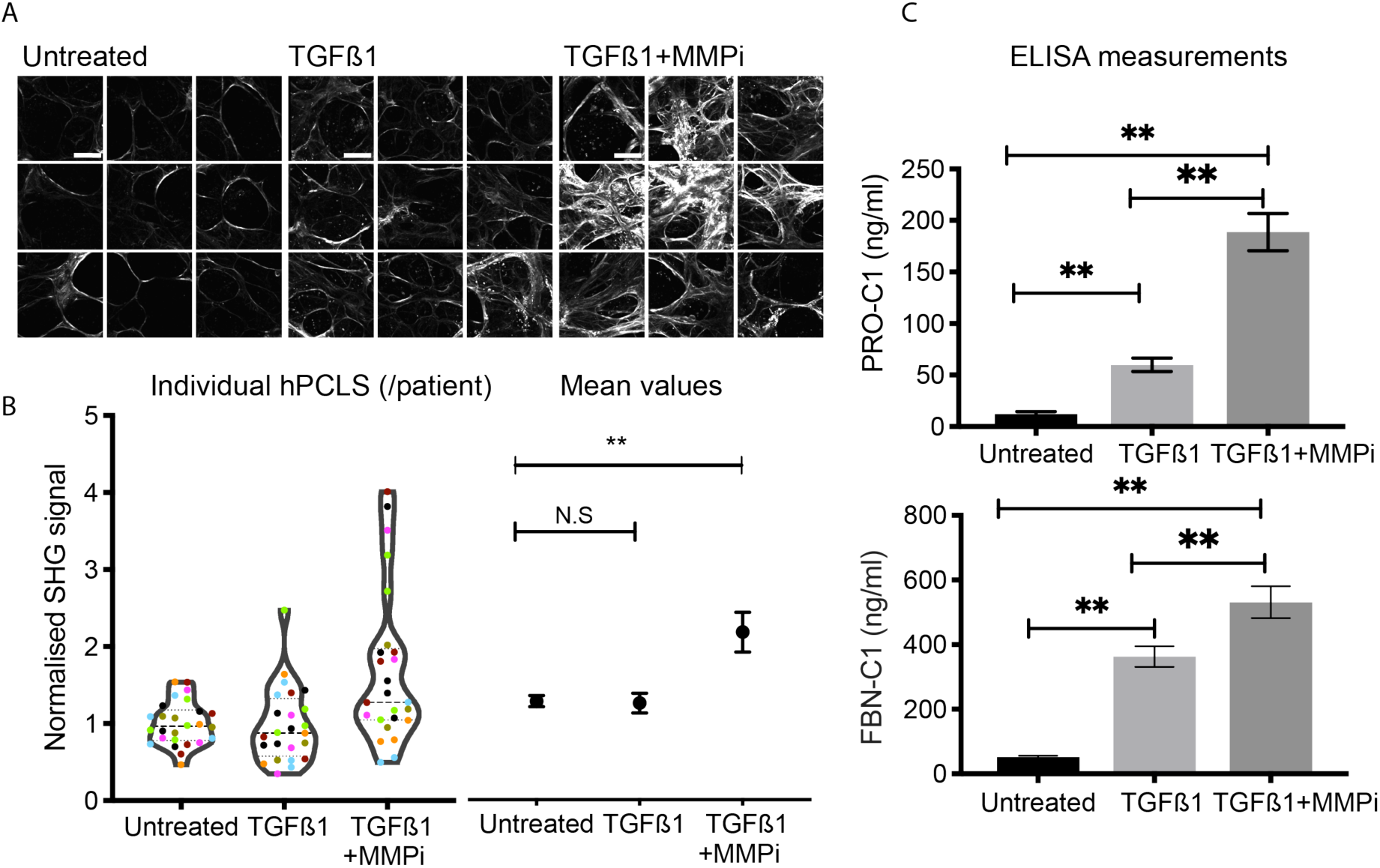
Second harmonic generation image analysis of fibrillar collagen deposition in ex vivo cultured hPCLS upon simultaneous stimulation with TGFß1 and metalloproteinase inhibitor (MMPI-GM6001): (A) Maximum Z-projection representative rois of vehicle, TGFß1 and TGFß1+MMPi (GM6001) treated hPCLS (same donor). Scale bar 250µm. (B) Individual as well as mean normalized SHG intensity signal value of hPCLS is shown. Students t-test (two-tailed) between means of different conditions was performed. Colored dots represent each hPCLS/ donor (7 donors with 3-4 hPCLS/ donor). Error bars represent SEM, p≤0.01. (C) ELISA measurement of soluble pro-fibrotic markers shows significantly more levels of both PRO-C1 and FBN-C upon concomitant TGFß1 and MMPi treatment compared to both vehicle and TGFß1 treated hPCLS. Error bars represent SEM. For ELISA data error bars SEM, *p<0.05, **p< 0.01***p<0.001, ANOVA with Turkey’s multiple comparisons test.

## Discussion

Interstitial lung diseases (ILDs) affect functional alveolar units within the lung parenchyma and are associated with high morbidity and mortality^51^. Animals studies and evaluation of IPF patient lung biopsies, peripheral blood proteome and bronchioalveolar lavage (BAL) have shown that TGFß1 plays a significant role in disease pathogenesis. Age^52^, smoking^53^ and MUC5 genetic variants^12^ are risk factors identified for IPF. The impact of cellular ageing on the etiology of pulmonary fibrosis, in particular, is difficult to study using rodent models and short time *in vitro* fibroblast cultures^54^. hPCLS model has emerged as a putative model system with high potential for recapitulating pathophysiological conditions close to that of human physiology. However, a characterization of the inherent molecular changes that such a system undergoes in *ex vivo* culture have not been reported to date.

In this study, we report for the first time proteomic and metabolomic changes of the hPCLS culture over two of ex vivo culture. Importantly we report that markers of the four major cell types are present after two weeks of *ex vivo* culture (Supplementary Figure 01C). Wound healing response has 4 phases^55^, involving an inflammatory phase of ECM degradation, and final proliferative ECM formation and remodeling phase^55^. Cell type-specific markers data suggest a persistent ECM degradative inflammatory phase present during the 2 weeks ex vivo culture. Indeed, we observe that hPCLSs have high immune cell markers (Supplementary Figure 01C) therefore, upregulated inflammatory pathways (Supplementary Figure 01B), sustained upregulation of pathways related to ECM degradation (Supplementary Fig 01B), downregulation of key structural ECM collagens (TableS2) and upregulation of proteins involved in fibrillar collagen degradation (TableS1). Our proteomics data also shows that hPCLSs are adapting to lack of serum in the culture media by upregulating cholesterol biosynthesis^56^, a process vital for the viability of biological membranes.

Interestingly, in contrast to the proteomic data, the metabolomic data of the hPCLS showed minor changes after the initial adaptation period (4 days).

On the other hand, a persistent inflammatory process together with the presence of other cell types such as epithelial, endothelial and immune cells, suggest a possibility of transition of these cell types into myofibroblasts upon stimulation with cytokines such as TGFß1^3^, hence eliciting a physiologically relevant pro-fibrotic response. Indeed, after treatment with TGFß1, ELISA measurements of soluble biomarkers^28^ of fibrosis such as PRO-C1 and FBN-C1 showed a substantial increase. Interestingly, repetitive stimulation resulted in sustained increased production of PRO-C1 and Fibronectin (Figure 2A) over time, showing no response attenuation to TGFß1. To get an in-depth knowledge of the nature of this pro-fibrotic response, we carried out the proteomic analysis of *ex vivo* cultured hPCLS (day 13, ±TGFß1 treatment). Consistent with other studies^10^ the analysis showed upregulation of many pro-fibrotic markers (Figure 2B) including pro-fibrotic fibrillar collagens COLI, COLIII and COLV. On day 13 of hPCLS ex vivo culture, a total of 109 proteins were significantly upregulated. Strikingly, 64 out of 65 upregulated proteins belong to mesenchymal cells and 40 out of 44 downregulated genes belong to epithelial cells (Supplementary Figure 1C). These data suggest that the pro-fibrotic signal induced after TGFß1 treatment of hPCLS is due to epithelial to mesenchymal transition^3^.

To further investigate the functional consequences of highly upregulated profibrotic response we sought to measure the deposition of fibrillar collagen (tissue phenotypic marker of fibrosis)^3^ in ECM of hPCLS. Other studies^10,20^ have made attempts to use hPCLS for studying fibrotic signalling, however, a functional readout that confirms excess extracellular matrix deposition has been lacking. Not only in this model system but in other systems as well, only indirect readouts such as ELISA^57^ of pro-collagen peptides or hydroxyproline levels in serum have been used to infer on fibrosis, e.g. in liver, kidney or lung^58–60^. In hPCLS model system, attempts have been made to quantify the deposition of fibrillar collagen using immunohistochemistry albeit qualitative^20^, therefore we applied label-free second harmonic imaging to specifically quantify fibrillar collagen content present in the ECM of hPCLS. A key advantage of label-free imaging is its ability to image deep into the tissues and specifically measure fibrillar ECM deposited collagen. The SHG image analysis of healthy and IPF derived hPCLS showed higher levels of fibrillar collagen in IPF hPCLS (Figure 3B-C), showing the quantitative nature of the established imaging workflow. Interestingly, upon SHG microscopy the increase in expression of pro-fibrotic markers, in particular, COLI (PRO-C1-ELISA, COL1A2-mass spec) did not result in a corresponding significant higher deposition in ECM of TGFß1 stimulated hPCLS (Figure 3E). Given hPCLS show upregulated expression of MMPs (steady-state ex vivo culture, Table S1), we analysed the Log2FC levels of MMPs in TGFß1 treated hPCLS proteomics (day 13). Indeed both MMP2 and MMP14 (MMPs responsible for degradation of fibrillar collagens)^24^ show an upregulated expression with Log2FC of 0.68 (fdr 0.16) and 0.52 (fdr 0.5) respectively. To enable the deposition process of fibrillar collagens in hPCLS, we tested whether the process of ECM degradation could be reduced by concomitantly treating the *ex vivo* cultured hPCLS with TGFß1 and the MMP inhibitor GM6001, Indeed, the media supernatants of hPCLS treated with TGFß1 and GM6001 have higher levels of PRO-C1 and Fibronectin levels compared to just TGFß1 stimulation (Figure 4C). Pulmonary fibrosis is essentially an ageing disorder^61^developing over several years and to recapitulate the hallmark in 2 weeks of culture time represent a real challenge. Here we show that concomitant treatment with TGFß1 and GM6001, could indeed enable the measurement of fibrils deposition in this *ex-vivo* model system.

The observation that the expression of soluble markers does not correlate to the deposition of fibrillar collagen in ECM further strengthens the need of an integrated workflow to measure both soluble and ECM deposited fibrils.

Furthermore, our SHG data analysis has highlighted the variability of inherent fibrillar collagen content in hPCLS (Supplementary Figure 03), showing the need for image analysis of each hPCLS, where only the selected areas are quantified, excluding SHG-confounding regions such as blood vessels and airways.

It should also be noted that not all patients respond equally to TGFß1+GM6001 (Figure 4B), there are hPCLS (donor-specific) that do not respond with corresponding increase in ECM deposition (Fig 4B). The reasons for this selective lack of deposition in certain hPCLS being donor-specific is hard to reason, however, IPF lungs develop fibrotic foci^62^, rather than diffuse fibrosis, one explanation could be that certain hPCLS are more predisposed for ECM deposition than others. These fibrotic foci have been characterized well and Jones et al.,^62^ have made a significant effort in this direction. Similar to this study, our analysis also focusses on interstitium of lung parenchyma and could in future help to identify the progression and development of such foci.

We propose that this method together with measurement of soluble markers such as PRO-C1, FBN-C, C3M could enable the characterization of the dynamics of collagen synthesis and degradation (the fractional synthesis) *ex vivo* upon fibrosis progression.

Our study provides an integrated in-depth characterization of the here presented hPCLS model system but it is important to note that there are additional points that could be further investigated in future studies, for example, different culture conditions such as microfluidic bioreactors or cell culture media composition. Our current imaging set-up enables a deep view into the tissue and live imaging of the fibrillar collagen dynamics would be the next frontier. Live SHG imaging can quantify disease progression as well as regression upon therapeutic intervention. It would be exciting to culture IPF derived hPCLS and use anti-fibrotic agents to assess over time the effects on fibrillar collagen. Furthermore, recently, hPCLS and non-human derived PCLS have also been subjected to siRNA knockdowns^63^ and viral transductions^64^, indicating the amenability of this model also to genetic manipulation for functional genomic studies.

## Supporting information

excel sheet-SuplementaryCellTypeComp-TimeCourse

excel sheet-Suplementarymetabolome

excel sheet SuplementaryProtTGF

excel sheet-SuplementaryProtTimeCourse

ImageAnalysis

## Acknowledgements

Tissue samples were provided by Lung Biobank Heidelberg, member of the Biomaterial Bank Heidelberg (BMBH) and the biobank platform of the German Center for Lung Research (DZL). Technical assistance of Christa Stolp in tissue assembling is gratefully acknowledged. We would also like to acknowledge the help of Viki Barret (GSK Stevenage) for help in training with Krumdieck. We sincerely thank Katrin Strohmer and Anna Rutkowska-Klute (Cellzome-GSK) for their help in setting up the SHG imaging. Pepperkok team is also acknowledged for their help with manuscript preparation and fruitful discussions for data analysis.

## Supplementary figures

**Supplementary Figure 01:**
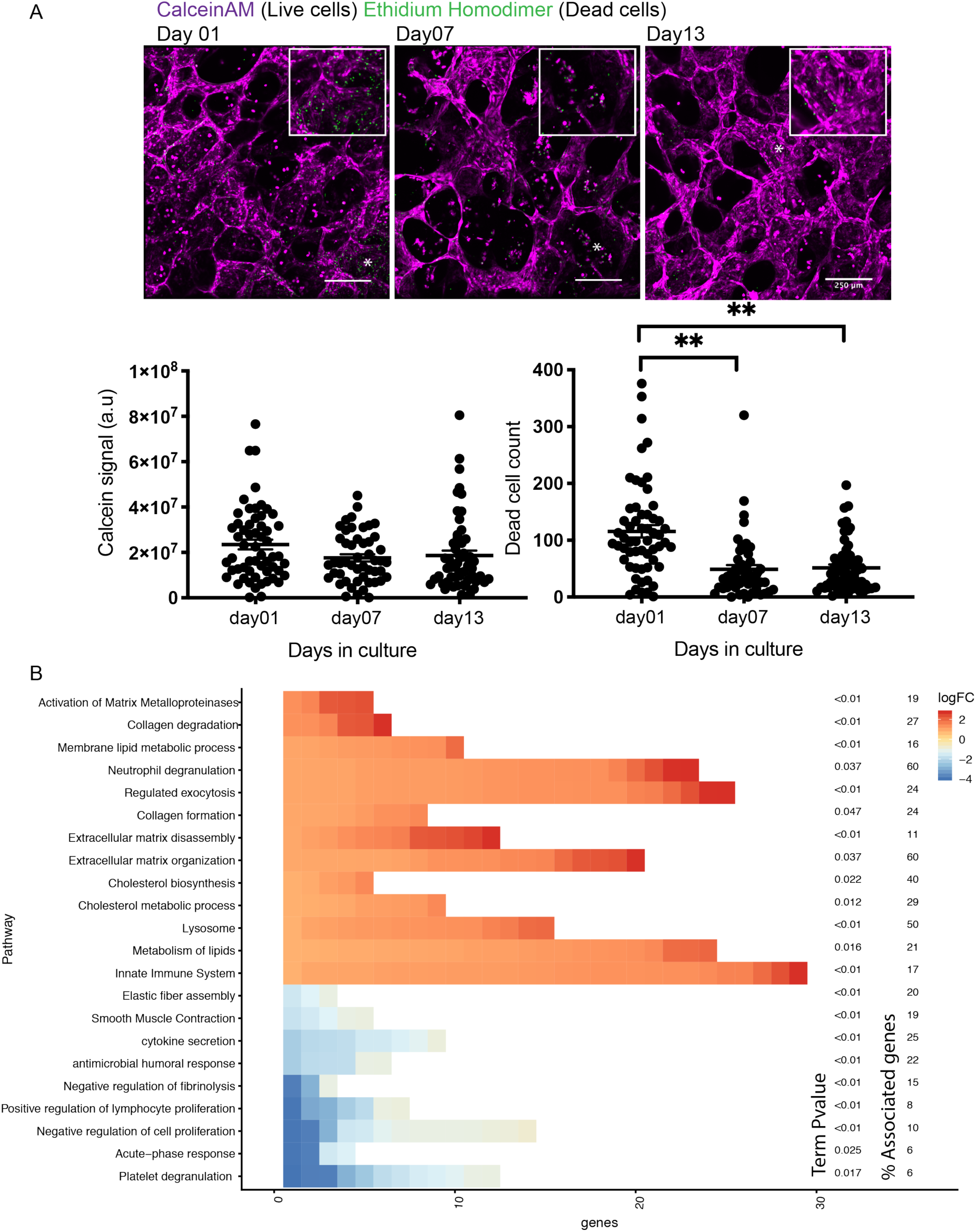
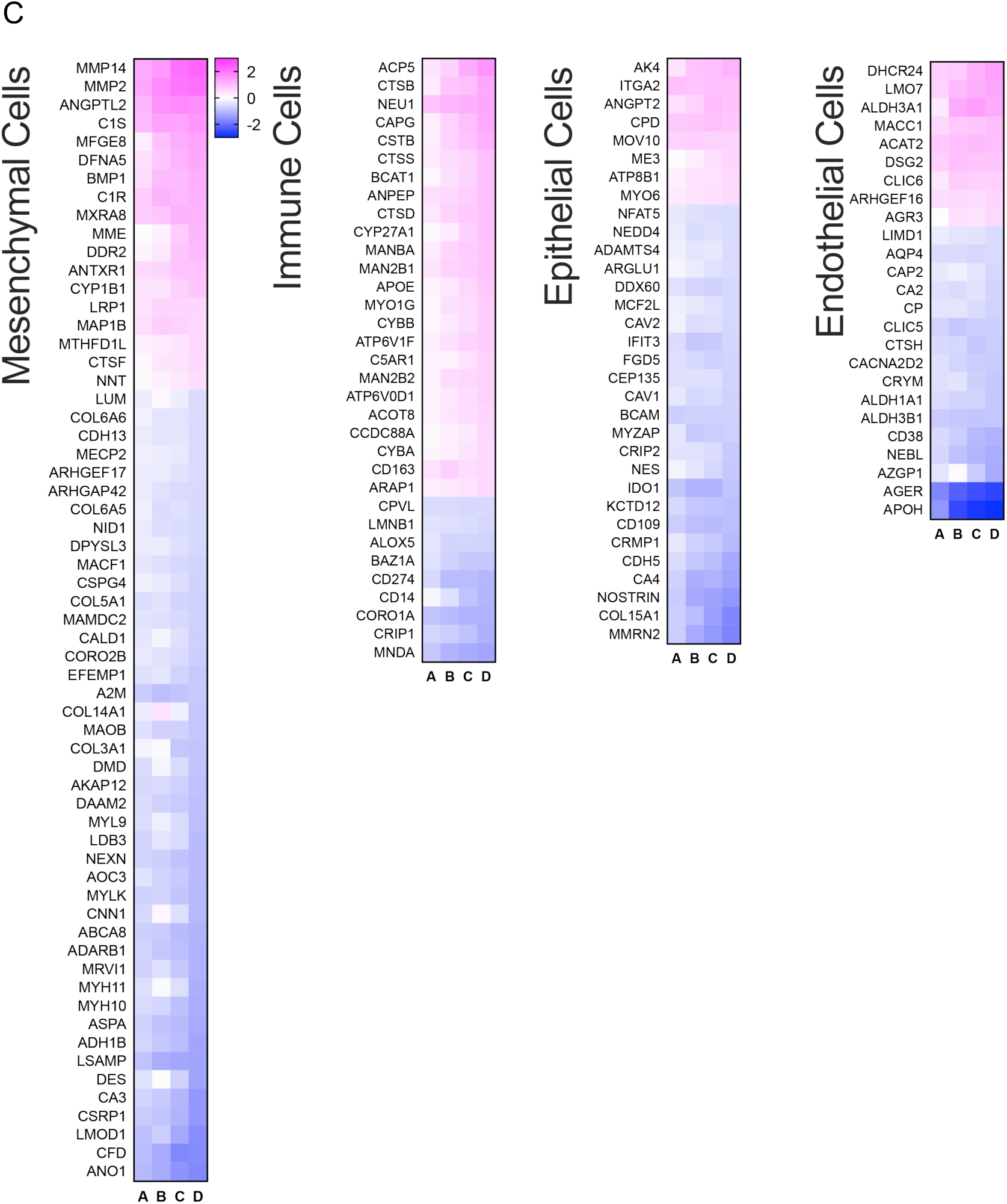
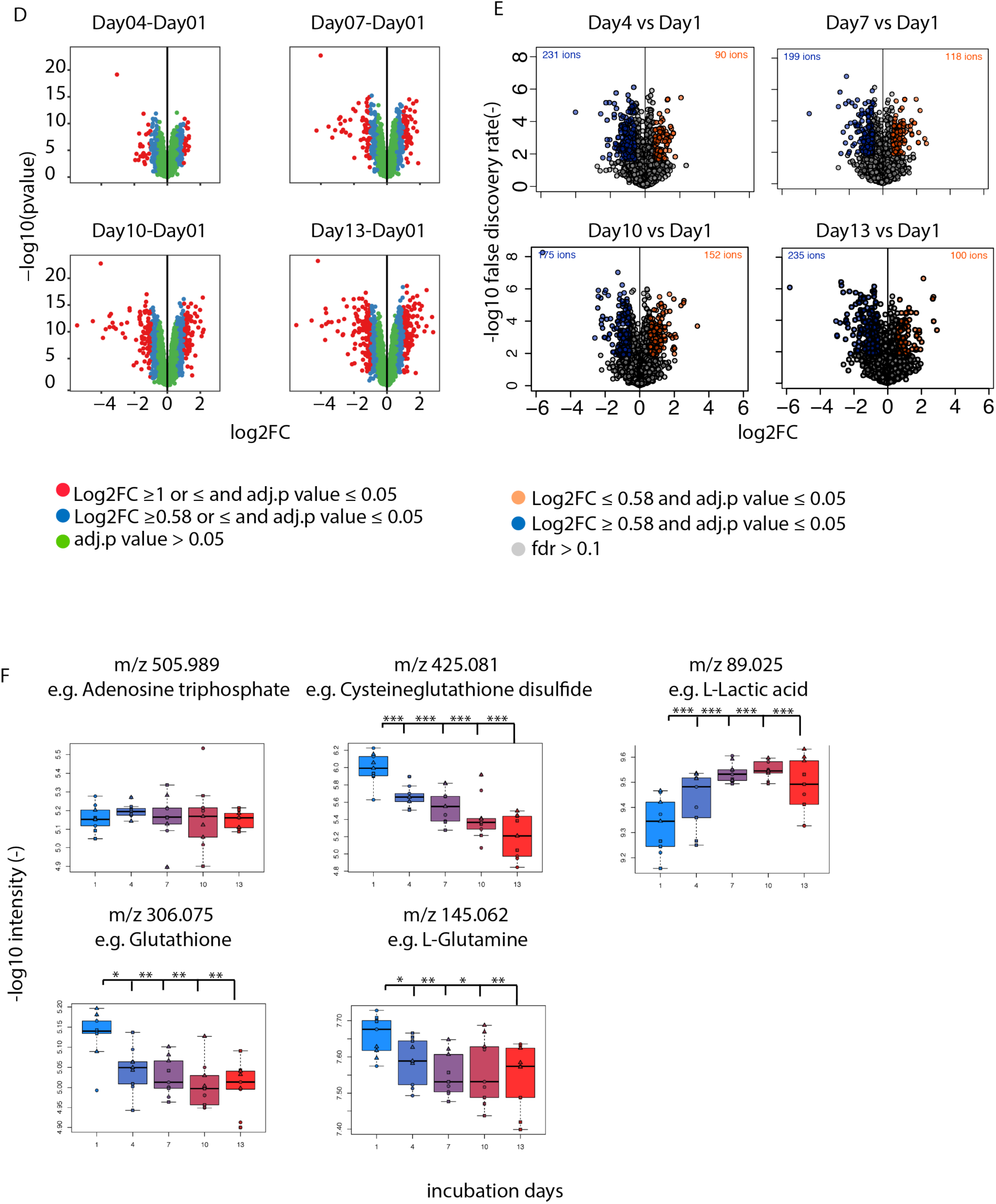
(A) Representative live confocal images of hPCLS incubated with Calcein AM (labels live cells) and Ethidium homodimer (Dead cells) and quantification live tissue (Calcein signal) and dead cell count (ethidium homodimer) Error bars represent SEM, and p-values were calculated using ordinary one-way ANOVA, **p<0.001 (B) Significantly regulated genes on day 13 compared to day 1 were subjected to Cytoscape pathway analysis. The number of genes in each enriched GO terms present in our data set was plotted. Term Pvalue represents the pvalue for association of these genes to whole of GO term. % associated genes, is the percentage of our genes overlapping with all of the members of a given GO term. (C) Significantly regulated genes (day 13 vs. Day 1) were compared with unique cell type markers as established in LGEA^23^ study. A, B, C and D mark comparisons of day4 vs day1, day7 vs day1, day10 vs day1 and day13 vs day1 respectively. (D) Volcano plots show Log2FC of all the genes detected against their respective fdr. Red dots represent highly significant genes, while as blue are those regulated that are significant but moderate in the level of change. (E) Metabolites detected upon LC/MS were similarly plotted as in (D). (F) Represent individual metabolites and their change over days in *ex vivo* culture.

**Table S1:** Log2FC of metalloproteinases across different days of *ex vivo* culture.

| gene_name | logFC-Day04 -Day 1 | fdr | logFC<br>Day 7-Day 1 | fdr | logFC<br>Day 10 -Day 1 | fdr | logFC<br>Day 13 - Day 1 | fdr |
| --- | --- | --- | --- | --- | --- | --- | --- | --- |
| MMP3 | 1.3 | 0.00 | 0.6 | 0.02 | 0.4 | 0.10 | 0.1 | 0.82 |
| **MMP2** | **1.3** | **0.00** | **1.7** | **0.00** | **2.0** | **0.00** | **2.1** | **0.00** |
| **MMP14** | **1.3** | **0.00** | **1.5** | **0.00** | **2.1** | **0.00** | **2.3** | **0.00** |
| **MMP1** | **1.1** | **0.00** | **1.0** | **0.00** | **1.2** | **0.00** | **1.3** | **0.00** |
| MMP10 | 0.4 | 0.17 | 0.6 | 0.02 | 0.5 | 0.05 | -0.1 | 0.82 |
| MMP12 | -0.1 | 0.88 | 1.2 | 0.00 | 1.7 | 0.00 | 2.8 | 0.00 |
| MMP28 | -0.1 | 0.61 | -0.2 | 0.43 | -0.3 | 0.19 | -0.3 | 0.22 |
| MMP9 | -0.2 | 0.51 | 0.3 | 0.36 | 0.5 | 0.05 | 1.2 | 0.00 |
| MMP19 | -0.2 | 0.05 | -0.2 | 0.02 | -0.2 | 0.05 | -0.3 | 0.00 |

**TableS2:** Log2FC of collagens across different days of *ex vivo* culture.

| gene_name | logFC-Day04 -Day 1 | fdr | logFC<br>Day 7-Day 1 | fdr | logFC<br>Day 10 -Day 1 | fdr | logFC<br>Day 13 - Day 1 | fdr |
| --- | --- | --- | --- | --- | --- | --- | --- | --- |
| COL8A1 | 0.0 | 0.99 | 0.3 | 0.46 | 0.1 | 0.89 | 0.1 | 0.73 |
| COL7A1 | 0.3 | 0.09 | 0.1 | 0.61 | -0.1 | 0.65 | -0.2 | 0.31 |
| COL6A3 | -0.1 | 0.57 | 0.1 | 0.57 | -0.1 | 0.60 | -0.2 | 0.07 |
| COL4A3 | -0.5 | 0.39 | -0.4 | 0.46 | -0.6 | 0.23 | -0.2 | 0.68 |
| COL5A2 | -0.1 | 0.64 | 0.1 | 0.55 | -0.2 | 0.47 | -0.2 | 0.30 |
| COL1A2 | 0.0 | 0.97 | 0.4 | 0.10 | -0.1 | 0.72 | -0.2 | 0.39 |
| COL16A1 | -0.2 | 0.54 | 0.0 | 0.91 | -0.3 | 0.39 | -0.3 | 0.48 |
| COL6A1 | -0.2 | 0.23 | -0.1 | 0.38 | -0.2 | 0.09 | -0.4 | 0.01 |
| COL6A2 | -0.1 | 0.29 | -0.1 | 0.38 | -0.2 | 0.12 | -0.4 | 0.01 |
| COL4A2 | -0.4 | 0.27 | -0.3 | 0.48 | -0.5 | 0.15 | -0.4 | 0.29 |
| COL1A1 | -0.1 | 0.71 | 0.3 | 0.32 | -0.3 | 0.39 | -0.4 | 0.17 |
| COL12A1 | -0.3 | 0.03 | -0.3 | 0.01 | -0.4 | 0.00 | -0.4 | 0.00 |
| COL6A6 | -0.2 | 0.11 | -0.4 | 0.00 | -0.4 | 0.00 | -0.5 | 0.00 |
| COL6A5 | -0.2 | 0.09 | -0.5 | 0.00 | -0.4 | 0.00 | -0.6 | 0.00 |
| **COL5A1** | **-0.5** | **0.00** | **-0.4** | **0.01** | **-0.6** | **0.00** | **-0.7** | **0.00** |
| **COL14A1** | **-0.2** | **0.41** | **0.4** | **0.15** | **-0.2** | **0.45** | **-0.8** | **0.00** |
| **COL3A1** | **-0.2** | **0.63** | **-0.1** | **0.86** | **-0.8** | **0.01** | **-0.9** | **0.00** |
| **COL18A1** | **-0.4** | **0.01** | **-0.4** | **0.00** | **-0.7** | **0.00** | **-1.0** | **0.00** |
| **COL15A1** | **-0.7** | **0.00** | **-1.0** | **0.00** | **-1.4** | **0.00** | **-1.7** | **0.00** |

**Supplementary Figure 02:**
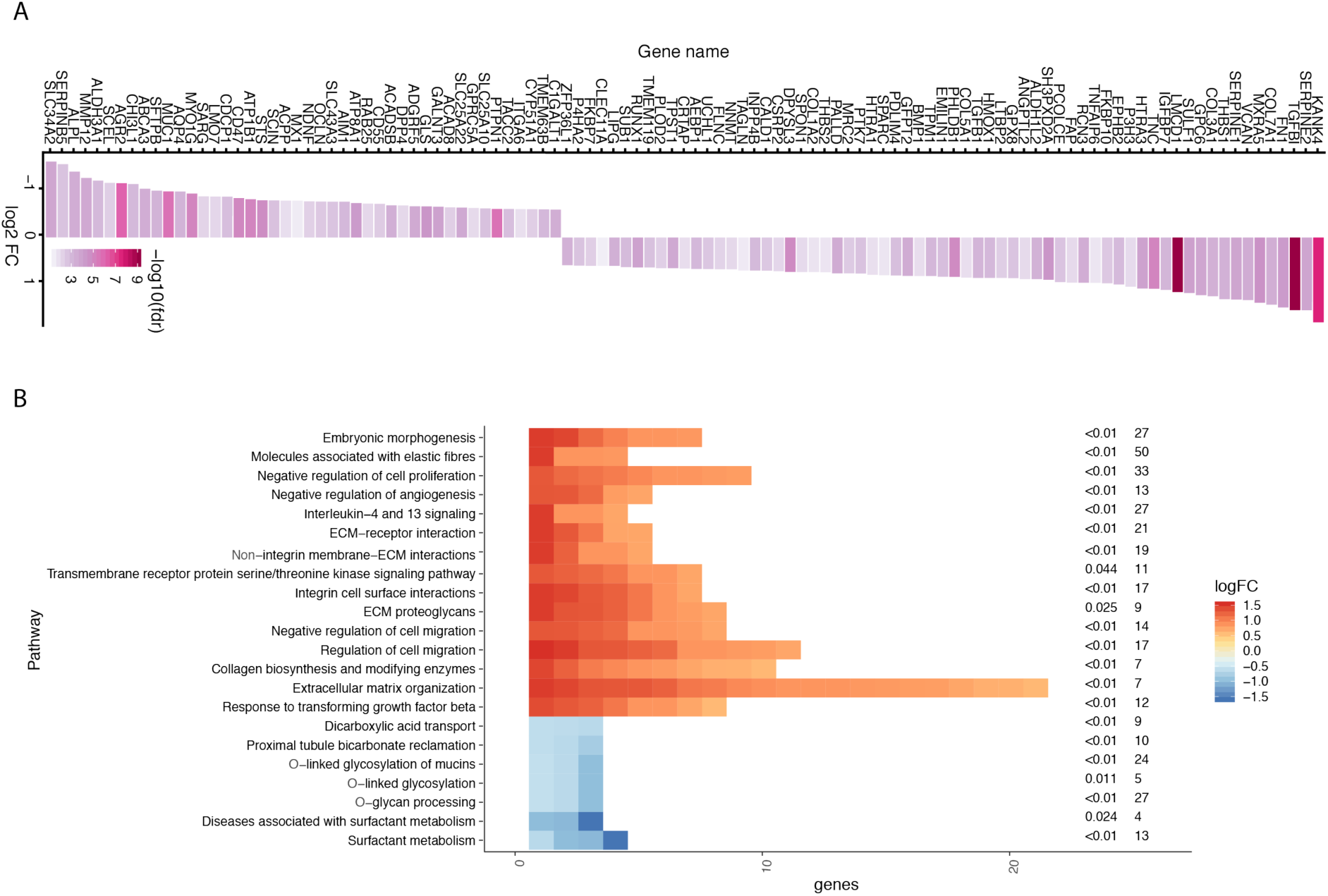
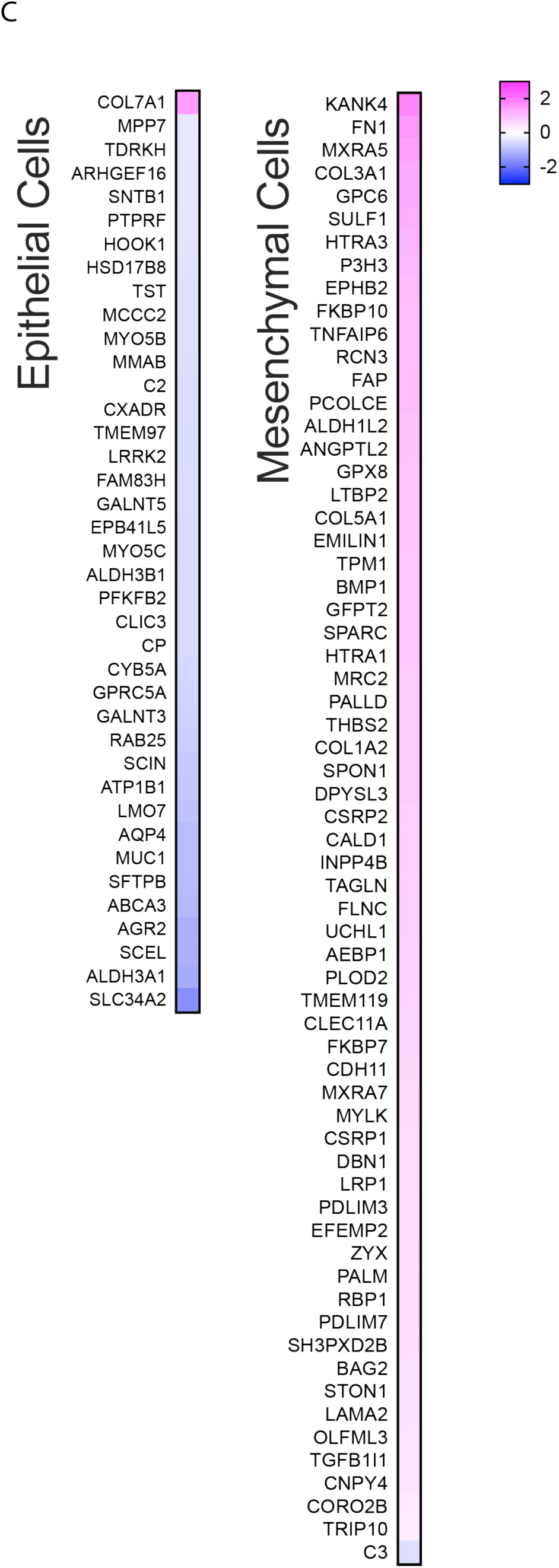
(A) List and level of significantly regulated genes upon TGFß1 treatment compared to untreated (day 13 vs. day 1) (B) Pathway analysis of significantly regulated genes upon TGFß1 treatment compared to untreated hPCLS (day 13 vs. day 1). The analysis was carried out using Cytoscape. (C) Significantly regulated genes were compared to different cell type markers as established by LGEA^23^ study and log2FC were plotted as a heat map.

**Supplementary Figure 03:**
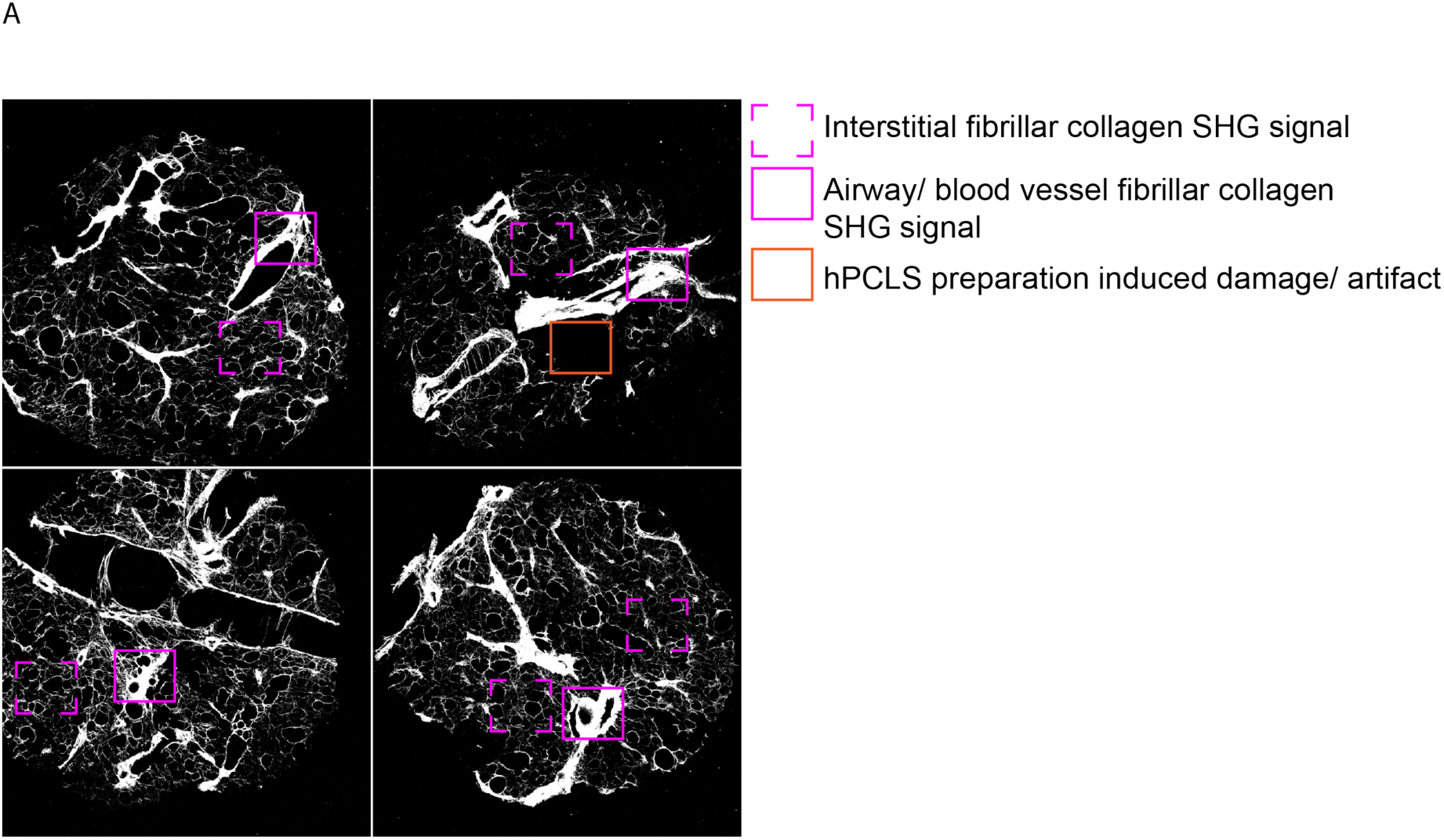
(A) Representative z-projection of SHG images from different hPCLS. Marked regions represent fibrillar collagen of different anatomical regions of hPCLS.

## Materials and methods

### Lung resection supply and licenses

Tumour-free human lung tissue resections from lung cancer patients were obtained from Thoraxklinik-Heidelberg with anonymized patient Ids. The patient consent and use of tissue was obtained as per the research ethics committee (Medical Faculty of University Heidelberg) approval reference number S-270/2001. Samples of ILD lung tissue were obtained from patients undergoing lung transplant for end–stage disease from Institute of Transplantation, Newcastle Upon Tyne Hospitals. The patient consent and use of ILD tissue was approved by national research ethics service (11/NE/0291) and UK Health Research Authority. The human biological samples were sourced ethically and their research use was in accord with the terms of the informed consents under an IRB/EC approved protocol.

### Precision-cut lung slice preparation and *ex vivo* culture

Healthy and IPF hPCLS were prepared as described previously^65^. Briefly, on the day when the tissue resection was received (day 0), lung tissue was inflated with 3% low melting agarose (Sigma#A9414) prepared in phenol free DMEM (Gibco# 41965-039). Next, 8 mm cores were prepared and 250 μm thick slices were generated using a Krumdieck tissue slicer. Tissue media was supplemented with penicillin, streptomycin, fungizone. The media (± stimulation) was replenished after every 72 hours till day 13. Cell culture grade plastic 24 well plates (Greiner#353226) were used to culture hPCLS. On day 0 media was additionally supplemented with 10% FCS (Gibco#10270106). The hPCLS were treated with 10ng/ml recombinant human TGFß 1 protein (R&DSystems#240-B-010) and 25µM (in 750µl of DMEM) of GM6001 (Merck#CC1010). Lived dead staining was carried out using calcein-am/ ethidium homodimer kit (ThermoFisher#L3224) as per supplier recommendations. The hPCLS were incubated in recommended concentration of live/dead reagents for 45 minutes prior to live confocal microscopy. hPCLS were always cultured in presence of penicillin-streptomycin (Gibco# 15140122) and amphotericin B (Giboc#15290026).

### ELISA measurements

Soluble extracellular matrix fragments (PINP, FBN-C, C3M) were determined in the media supernatant by competitive (ELISA) assays developed and validated by Nordic Bioscience A/S (Herlev, Denmark). According to the manufacturer’s instructions^29 66 67^. Values measured below the detection limit of the assay were assigned the lower limit of detection (LLOD).

### Mass spectrometry sample preparation and analysis

#### Proteomics

Briefly, hPCLS were washed with PBS and harvested using snap freezing in liquid Nitrogen. Upon thawing, samples (2 hPCLSs/technical replicate) were dissolved in 300μl of 2% SDS-H_2_O (with protease and phosphatase inhibitor) for 2 hours at room temperature. Subsequently, mechanically homogenized using bead ruptor. Before LC-MS/MS analysis, protein concentrations were measured (Pierce 660nm kit# 22660). Next samples were solubilized in 2 × SDS sample buffer and subjected to short SDS gel electrophoresis. Samples were further processed for LC-MS/MS analysis.

SDS PAGE gels were Coomassie stained. Gel lanes were cut into three slices covering the entire separation range (∼2 cm) and subjected to in-gel tryptic digestion^68^. Peptides were labeled via isobaric mass tags (TMT10, Thermo Fisher Scientific, Waltham, MA).TMT labeling was performed using the 10-plex TMT reagents, enabling relative quantification of 10 conditions in a single experiment^69^. Briefly, the labeling reaction was performed in 40 mM triethylammonium bicarbonate, pH 8.53 at 22°C and quenched with glycine. Labeled peptide extracts were combined to a single sample per experiment. Lyophilized samples were re-suspended in 1.25% ammonia in water and subjected to LC-MS/MS^69^.

Peptide and protein identification & identification Mascot 2.4 (Matrix Science, Boston, MA) was used for protein identification by using a 10 parts per million mass tolerance for peptide precursors and 20 mD (HCD) mass tolerance for fragment ions. The search database consisted of a customized version of the International Protein Index protein sequence database combined with a decoy version of this database created by using scripts supplied by MatrixScience.

Reporter ion intensities were read from raw data and multiplied with ion accumulation times (the unit is milliseconds) so as to yield a measure proportional to the number of ions; this measure is referred to as ion area ^70^. Spectra matching to peptides were filtered according to the following criteria: mascot ion score > 15, signal-to-background of the precursor ion > 4, and signal-to-interference > 0.5 ^70^. Fold changes were corrected for isotope purity as described and adjusted for interference caused by co-eluting nearly isobaric peaks as estimated by the signal-to-interference measure ^70^. Protein quantification was derived from individual spectra matching to distinct peptides by using a sum-based bootstrap algorithm; 95% confidence intervals were calculated for all protein fold changes that were quantified with more than three spectra ^70^.

### Metabolomics

hPCLSs were snap-frozen on the day of harvest. On the day of sample preparation, 2hPCLS/ technical replicate were shortly rinsed in 75mM Ammonium bicarbonate (pH 7.4) and mechanically homogenised in MS grade H_2_O to extract metabolites. Untargeted metabolomics analysis was performed as described^71^. Briefly, samples were analyzed on a LC/MS platform consisting of a Thermo Scientific Ultimate 3000 liquid chromatography system with autosampler temperature set to 10° C coupled to a Thermo Scientific Q-Exactive Plus Fourier transform mass spectrometer equipped with a heated electrospray ion source and operated in negative ionization mode. The isocratic flow rate was 150 μL/min of mobile phase consisting of 60:40% (v/v) isopropanol: water buffered with 1 mM ammonium fluoride at pH 9 and containing 10 nM taurocholic acid and 20 nM homotaurine as lock masses. Mass spectra were recorded in profile mode from 50 to 1,000 m/z with the following instrument settings: sheath gas, 35 a.u.; aux gas, 10 a.u.; aux gas heater, 200° C; sweep gas, 1 a.u.; spray voltage, −3 kV; capillary temperature, 250° C; S-lens RF level, 50 a.u; resolution, 70k @ 200 m/z; AGC target, 3×10^6^ ions, max. inject time, 120 ms; acquisition duration, 60 s. Spectral data processing was performed using an automated pipeline in R as described previously^71^. Detected ions were tentatively annotated as metabolites based on matching accurate mass within a tolerance of 5 mDa using the Human Metabolome database^72^.

### Data analysis

#### Proteomics

The R programming language (ISBN 3-900051-07-0) was employed to process the proteins output files of IsobarQuant. Only proteins which were quantified with at least two unique peptides and which were quantified in all experiments were used for further analysis. The “sumionarea_protein” columns were annotated to different experimental conditions. Then, potential batch-effects were removed using the limma package^73^ and the results were normalized using the vsn package^74^. Limma was also used to test for differential expression of proteins. A protein was considered significant with a 2-fold difference and a false discovery rate below 5 %. All significant proteins were clustered (hierarchical clustering – ward.d2 method) based on their Euclidean distances of log2 ratios towards the respective control.

### Network analysis using Cytoscape

Network analysis was carried out open source software platform Cytoscape^75^ was used. Briefly, genes that were significantly regulated (log2fc ≥ 0.58 or ≤ and adjusted p-value ≤0.05) were curated and subjected to pathway analysis. The graphs were plotted using an in-house R script.

### Second Harmonic generation imaging and sample preparation

On the day of harvest, the media supernatant was aspirated from each well containing hPCLS. hPCLS were washed with PBS (3X 1ml) and fixed in 1 ml of 4% PFA (EMS EM grade#5710) overnight at 4°C. Subsequently, hPCLS were washed with PBS (3X). Next, each PFA fixed hPCLS was transferred to 24 well plate with glass bottom (Greiner senso#662892). hPCLS were covered with 150µl of PBS and pressed to the bottom using a circular glass cover slip and plastic ring (made locally at EMBL workshop). Next, each well was further covered with 300µl of PBS to avoid drying of hPCLS (due to evaporation of PBS in the process of image acquisition). SHG imaging of hPCLS was performed on Zeiss NLO LSM780 microscope equipped with a pulsed multiphoton laser. hPCLS were excited with 2-photon wavelength of 880nm with a 20X (0.8 NA, air objective). 150-180µm deep z-stacks were acquired with a square tiled (7-8 × 7-8mm) scan. 2-photon excited signals originating from hPCLS were captured in 3 different channels backward SHG, forward SHG and 2-photon excited autofluorescence (2PEA). Forward SHG signal was captured using a laser blocking filter and a band pass filter 436/20 nm. The backward SHG signal was recorded using internal detectors in band width of 435-455 nm. 2PEA was detected in a bandwidth of 550-650 nm

### SHG image analysis

Image analysis of z-stacks consisting of SHG channels (backward and forward) and 2PEA were analysed using a semi-automated Jython based FIJI pipeline. Pipeline is available as supplementary document as well and is free for usage. Briefly, all the channels were thresholded, the area or the number of pixels under the threshold were counted for each channels (PBT), integrated intensity of PBT was also calculated for each channel (Sumintensity). To measure the total fibrillar collagen content, Sumintensity of PBT of backward and forward SHG signal were summed up to termed as total fibrillar collagen content. A projection of sumSHG (2D) image was created as well and this image was divided into 8×8 equally sized ROIs. And each ROI was manually classified as interstitial collagen or Non-interstitial collagen. Only ROIs of interstitial collagen were averaged per hPCLS SHG image. Each experiment has untreated & treated hPCLS SHG images. Sumintensity of total collagen content pixels/ ROI was normalized to the average SHG signal in untreated (average of all the ROIs) hPCLS of the respective donor. For each hPCLS SHG stack a minimum of 20 ROIs were analysed.

### FIJI code for interstitial and nor-interstitial ROI selection

Available as a compressed zip file (ImageAnalysis)

