## Supplementary material for "Multiomic and quantitative label-free microscopy-based analysis of *ex vivo* culture and TGFbeta1 stimulation of human precision-cut lung slices": ImageAnalysis: hPCLS-microscopy-image-analysis.py

```
from ij.gui import Roi,WaitForUserDialog
from ij.plugin import ZProjector, RGBStackMerge, SubstackMaker, Concatenator
from ij.plugin import Duplicator, ImageCalculator
from ij import IJ, ImagePlus, ImageStack, WindowManager,Prefs

from ij.process import StackStatistics

from ij.plugin.frame import RoiManager


from fiji.util.gui import GenericDialogPlus

import os.path, re, sys

from jarray import array


from loci.plugins import BF


from automic.table import TableModel            # this class stores the data for the table

from automic.table import ManualControlFrame    #this class visualises TableModel via GUI

from automic.parameters import ParameterCollection, ParameterType
from automic.parameters.xml import ParameterXmlWriter
from automic.utils import FileUtils
from automic.utils.roi import ROIManipulator2D

from java.io import File
from java.awt import Font
from java.lang import Integer


defDirectoryPrefsTag='tissue.analysis.default.directory'

def getGridRois(_image,_nFragmentsX, _nFragmentsY):
    width=_image.getWidth()
    height=_image.getHeight()
    fragmentWidth=width/_nFragmentsX
    fragmentHeight=height/_nFragmentsY
    gridRois=[]
    for indexX in range(0,_nFragmentsX):
        for indexY in range(0,_nFragmentsY):
            r=Roi(fragmentWidth*indexX,fragmentHeight*indexY,fragmentWidth,fragmentHeight)
            r.setName('%02d-%02d' %((indexY+1),(indexX+1)))
            gridRois.append(r)
    return gridRois

def filterRoisManually(_rawRois, _image):
    _image.show()
    
    IJ.setTool("hand") # "hand" tool seems to be needed for "Ctrl+mouse click" option to select ROIs in the ROI manager by clicking on the image
    
    roiManager=ROIManipulator2D.getEmptyRm()
    for r in _rawRois:
        roiManager.addRoi(r)
    roiManager.runCommand("UseNames", "true")
    roiManager.runCommand(_image,"Show All with labels");

    WaitForUserDialog('Manual action required','Remove unwanted grid Rois from ROI Manager').show()
    _image.hide()
    filteredRois=roiManager.getRoisAsArray()
    roiManager.runCommand("Reset")
    
    return filteredRois

def getCombinationsList(_nChannels):
    resultList=list()
    for f in range(1,_nChannels+1):
        for s in range(f+1,_nChannels+1):
            resultList.append((f,s))
    return resultList

def createFileColumnSubfolder(_table, _tag, _type):
    _table.addFileColumns(_tag,_type)
    FileUtils.createCleanFolder(_table.getRootPath(),_tag)

def createImageTableModel( _analysisPath, _nChannels):

    tbModel = TableModel(_analysisPath)

    tbModel.addColumn('DatasetId')
    tbModel.addFileColumns('RAW','IMG')

    # pixels above threshold (PBT)
    for i in range(1, _nChannels+1):
        tbModel.addValueColumn("PBT.%d" %i, "NUM")
    
    # total intensity in whole stack (no masking and no bg-subtraction)
    for i in range(1, _nChannels+1):
        tbModel.addValueColumn("SumIntensity.%d" %i, "NUM")
    
    # overlapping PBTs in different channels
    if _nChannels>1:
        indexes=getCombinationsList(_nChannels)
        for f,s in indexes:
            tbModel.addValueColumn("PBT.%dAND%d" %(f,s), "NUM")

    if _nChannels>2:
        tbModel.addValueColumn("PBT.1OR2OR3", "NUM")
    
    # MaxProjection for the sum of second harmonic channels
    createFileColumnSubfolder(tbModel, 'Second.Harmonic.Max.Proj','IMG')

    
    # binary and gated output images
    for i in range(1, _nChannels+1):
        createFileColumnSubfolder(tbModel, 'BW.%d' %i,'IMG')
        createFileColumnSubfolder(tbModel, 'GATED.%d' %i,'IMG')

    return tbModel

def createRegionTableModel( _analysisPath, _nChannels):
    tbModel = createImageTableModel( _analysisPath, _nChannels)
    createFileColumnSubfolder(tbModel, 'Region.Roi','ROI')
    
    tbModel.addValueColumn('Success','BOOL')
    return tbModel


def saveImageTable(_image,_fileName, _table, _iDataset, _tag):
    fullPath= os.path.join(_table.getRootPath(),_tag,_fileName)
    IJ.saveAs(_image, "Tiff", fullPath)
    _table.setFileAbsolutePath(fullPath, _iDataset, _tag, "IMG")


def close_all_image_windows():

    # forcefully closes all open images windows

    ids = WindowManager.getIDList();

    if (ids==None):

        return

    for i in ids:

        imp = WindowManager.getImage(i)

        if (imp!=None):

            win = imp.getWindow()

            if (win!=None):

                imp.changes = False # avoids the "save changes" dialog

                win.close()


def extractChannel(imp, nChannel, nFrame):

    """ Extract a stack for a specific color channel and time frame """

    stack = imp.getImageStack()

    ch = ImageStack(imp.width, imp.height)

    for i in range(1, imp.getNSlices() + 1):

        index = imp.getStackIndex(nChannel, i, nFrame)

        ch.addSlice(str(i), stack.getProcessor(index))

    return ImagePlus("Channel " + str(nChannel), ch)


def measureSumIntensity3D(imp):

    stats = StackStatistics(imp)

    return stats.mean * stats.pixelCount


def autoThreshold(imp, method):

    impout = imp.duplicate() 

    IJ.run(impout, "Auto Threshold", "method=" + method + " white stack use_stack_histogram");

    impout.setTitle("Auto Threshold")

    return impout


def threshold(_img, lower_threshold, upper_threshold):

    imp = Duplicator().run(_img)

    IJ.setThreshold(imp, lower_threshold, upper_threshold)

    IJ.run(imp, "Convert to Mask", "stack")

    imp.setTitle("Threshold");

    return imp


def getChannelMaxProgection(_originalImage,_channelIndex):
    #extract channel image
    # WARNING: image copy is not created here
    channelImage=extractChannel(_originalImage, _channelIndex, 1)#Duplicator().run(_originalImage,_segmentationChannelIndex,_segmentationChannelIndex,1,imp.getNSlices(),1,1)
    
    #project Stack
    projector=ZProjector()
    projector.setMethod(ZProjector.MAX_METHOD)
    projector.setImage(channelImage)
    projector.doProjection()
    projectedImage=projector.getProjection()
    projectedImage.setTitle('Channel MAX projection')
    
    return projectedImage


def addRowToRegionTable(_regionTable, _imageTable, _imageTableRowIndex):
    _regionTable.addRow()
    regionTableRowIndex = _regionTable.getRowCount()-1
    imageTableColumnNames = _imageTable.getColumnNames()
    for columnName in imageTableColumnNames:
        _regionTable.setValueAt(_imageTable.getValueAt(_imageTableRowIndex,columnName),regionTableRowIndex,_regionTable.getColumnIndex(columnName))
    _regionTable.setBooleanValue(True,regionTableRowIndex,'Success')
    return regionTableRowIndex


def analyze(iDataSet, tbModel, tbModelRegions, _analysisParameters):

    

    #

    # LOAD FILES

    #

    

    filepath = tbModel.getFileAbsolutePathString(iDataSet, "RAW", "IMG")

    filename = tbModel.getFileName(iDataSet, "RAW", "IMG")
    basename=os.path.splitext(filename)[0]
    filenametif = '%s.tif' %(basename)
    
    print("Analyzing: "+filepath)


    nChannels=_analysisParameters.getParameterValue ("Number of Channels")
    
    originalImage=BF.openImagePlus(filepath)[0]
    if originalImage.getNChannels()!=nChannels:
        raise Exception('Incompatible number of channels in the dataset %s' %filename)
    
    
    IJ.run("Options...", "iterations=1 count=1"); 

    

    #Create image which is a sum of max projections of second harmonic images
    segStack=ImageStack(originalImage.getWidth(),originalImage.getHeight())
    for channelIndex in _analysisParameters.getParameterValue("Segmentation Channels"):
        segStack.addSlice(getChannelMaxProgection(originalImage,channelIndex).getProcessor())
    sProjector=ZProjector()
    sProjector.setMethod(ZProjector.SUM_METHOD)
    sProjector.setImage(ImagePlus('Stack of Max projections',segStack))
    sProjector.doProjection()
    sumSegMaxProjectionImage=sProjector.getProjection()
    output_filename = "MaxProj--%s" %filenametif
    saveImageTable(sumSegMaxProjectionImage,output_filename, tbModel, iDataSet, 'Second.Harmonic.Max.Proj')
    sumSegMaxProjectionImage.setTitle('Sum of max projections for segmentation channels')
    
    
    #create lines in Regions table and copy data there
    regionIndexRoi=[]
    gridRois=getGridRois(originalImage,_analysisParameters.getParameterValue("Region Split X"), _analysisParameters.getParameterValue("Region Split Y"))

    if _analysisParameters.getParameterValue('Filter grid regions manually'):
        gridRois=filterRoisManually(_rawRois=gridRois, _image=sumSegMaxProjectionImage)
    
    regionRoiLocation = os.path.join(tbModelRegions.getRootPath(),'Region.Roi')
    
    for gridRoi in gridRois:
        regionTableRowIndex = addRowToRegionTable(_regionTable = tbModelRegions, _imageTable = tbModel, _imageTableRowIndex = iDataSet)
        regionIndexRoi.append((regionTableRowIndex,gridRoi))

        regionRoiFileName = '%s--%s' %(basename,gridRoi.getName())
        ROIManipulator2D.saveRoiToFile(regionRoiLocation,regionRoiFileName,gridRoi)
        tbModelRegions.setFileAbsolutePath(regionRoiLocation,'%s.zip' %regionRoiFileName,regionTableRowIndex,'Region.Roi','ROI')
        pass
    
    
    impA = []

    sumIntensities = []

    sumIntensitiesBW = []

    

    for iChannel in range(1,nChannels+1):

        
        # gating

        lower_threshold_value = _analysisParameters.getParameterValue("LOWER_TH_CH%d" %iChannel)

        upper_threshold_value = _analysisParameters.getParameterValue("UPPER_TH_CH%d" %iChannel)
        print "gating channel %d with values from %d to %d" %(iChannel,lower_threshold_value,upper_threshold_value)
        imp_c = extractChannel(originalImage, iChannel, 1)

        imp_bw = threshold(imp_c, lower_threshold_value, upper_threshold_value) 

        impA.append(imp_bw)  # store the binary mask (0, 255) for OR operations later

        
        # save binary image
        output_filename = "BW.Ch%d--%s" %(iChannel,filenametif)
        saveImageTable(imp_bw,output_filename, tbModel, iDataSet, "BW.%d" %iChannel)
        for regionIndex,regionRoi in regionIndexRoi:
            tbModelRegions.setFileAbsolutePath(os.path.join (output_folder,"BW.%d" %iChannel), output_filename, regionIndex, "BW.%d" %iChannel, "IMG")
        
        # convert from 0,255 to 0,1 for the following calculations

        imp_01 = imp_bw.duplicate()

        IJ.run(imp_01, "Divide...", "value=255 stack"); 

        

        # set all pixels in the intensity image to zero that are not within the gate

        imp_c_gated = ImageCalculator().run("Multiply create stack", imp_c, imp_01)

    

        # save gated image
        output_filename = "GATED.Ch%d--%s" %(iChannel,filenametif)
        saveImageTable(imp_c_gated,output_filename, tbModel, iDataSet, "GATED.%d" %iChannel)
        for regionIndex,regionRoi in regionIndexRoi:
            tbModelRegions.setFileAbsolutePath(os.path.join (output_folder,"GATED.%d" %iChannel), output_filename, regionIndex, "GATED.%d" %iChannel, "IMG")

        
        # now measure in imp_c_gated and imp_bw(0,1)

        tbModel.setNumericValue(round(measureSumIntensity3D(imp_01),0), iDataSet, "PBT.%d" %iChannel)

        tbModel.setNumericValue(round(measureSumIntensity3D(imp_c_gated),0), iDataSet, "SumIntensity.%d" %iChannel)


        for regionIndex,regionRoi in regionIndexRoi:
            imp_01.setRoi(regionRoi)
            imp_c_gated.setRoi(regionRoi)
            tbModelRegions.setNumericValue(round(measureSumIntensity3D(imp_01),0), regionIndex, "PBT.%d" %iChannel)
            tbModelRegions.setNumericValue(round(measureSumIntensity3D(imp_c_gated),0), regionIndex, "SumIntensity.%d" %iChannel)
        
        imp_01.setRoi(None)
        imp_c_gated.setRoi(None)
    

    # Compute overlaps of binary images

    if nChannels>1:
        indexes=getCombinationsList(nChannels)
        for f,s in indexes:
            imp_bw = ImageCalculator().run("AND create stack", impA[f-1], impA[s-1])
            tbModel.setNumericValue(measureSumIntensity3D(imp_bw)/255, iDataSet, "PBT.%dAND%d" %(f,s))
            
            for regionIndex,regionRoi in regionIndexRoi:
                imp_bw.setRoi(regionRoi)
                tbModelRegions.setNumericValue(measureSumIntensity3D(imp_bw)/255, regionIndex, "PBT.%dAND%d" %(f,s))
        
        imp_bw.setRoi(None)
    


    
    # Compute total volume, i.e. pixels that are above threshold in any of the channels

    if nChannels>2:
        imp_combined = ImageCalculator().run("OR create stack", impA[0], impA[1])

        imp_combined = ImageCalculator().run("OR create stack", imp_combined, impA[2])

        tbModel.setNumericValue(round(measureSumIntensity3D(imp_combined)/255,0), iDataSet, "PBT.1OR2OR3")

        
        for regionIndex,regionRoi in regionIndexRoi:
            imp_combined.setRoi(regionRoi)
            tbModelRegions.setNumericValue(round(measureSumIntensity3D(imp_combined)/255,0), regionIndex, "PBT.1OR2OR3")
        
        imp_combined.setRoi(None)

#

# ANALYZE INPUT FILES

#

def determine_input_files(foldername, tbModel, _extension):

    

    print("Determine input files in:",foldername)

    PBTtern = re.compile('(.*).'+_extension) 

    

    i = 0

    for root, directories, filenames in os.walk(foldername):

        for filename in filenames:

            print("Checking:", filename)

            match = re.search(PBTtern, filename)

            if (match == None) or (match.group(1) == None):

                continue

            tbModel.addRow()

            tbModel.setFileAbsolutePath(foldername, filename, i, "RAW","IMG")

            tbModel.setStringValue(filename, i,'DatasetId')
            print("Accepted:", filename)

            
            i += 1

    

    return tbModel


#

# GET PARAMETERS

#

def get_parameters():
    
    inputCollection=ParameterCollection()
    
    
    headerFont=Font(Font.SERIF,Font.BOLD,14)
    
    gd = GenericDialogPlus("Enter analysis settings")
    gd.addMessage("Input Data", headerFont)
    

    gd.addStringField("Datasets to Analyse", "all");
    gd.addStringField("Input File extension", "czi");
    gd.addNumericField("Number of Channels",4,0)
    gd.addDirectoryField("Input Data Path", Prefs.get(defDirectoryPrefsTag,None),40)
    
    gd.addMessage("Options to exclude regions from quantification",headerFont)
    
    gd.addCheckbox('Filter grid regions manually',True)
    gd.addNumericField('Region Split X',3,0)
    gd.addNumericField('Region Split Y',3,0)
    
    
    
    gd.showDialog()
    if gd.wasCanceled():
        WaitForUserDialog("Analysis aborted by User").show()
        sys.exit("Analysis aborted by User")
    
    inputCollection.addParameter("Datasets to Analyse",gd.getNextString(),None,ParameterType.STRING_PARAMETER)
    inputCollection.addParameter("Input File Extension",gd.getNextString(),None,ParameterType.STRING_PARAMETER)
    inputCollection.addParameter("Number of Channels",int(gd.getNextNumber()),None,ParameterType.INT_PARAMETER)
    inputCollection.addParameter("Input Data Path",gd.getNextString(),None,ParameterType.FOLDERPATH_PARAMETER)
    Prefs.set(defDirectoryPrefsTag, inputCollection.getParameterValue("Input Data Path"))
    
    inputCollection.addParameter("Filter grid regions manually",gd.getNextBoolean(),None,ParameterType.BOOL_PARAMETER)
    inputCollection.addParameter("Region Split X",int(gd.getNextNumber()),None,ParameterType.INT_PARAMETER)
    inputCollection.addParameter("Region Split Y",int(gd.getNextNumber()),None,ParameterType.INT_PARAMETER)
    
    
    
    nChannels = inputCollection.getParameterValue ("Number of Channels")
    
    gd = GenericDialogPlus("Specify Channels and Thresholds")
    gd.addMessage("Channel thresholds",headerFont)
    gd.addMessage("Please note: the threshold values are inclusive!\nThus, to exlude pixels with value 255 the upper threshold needs to be 254")
    
    
    for chIndex in range (1,nChannels+1):
        gd.addNumericField('LOWER_TH_CH%d' %chIndex, 30, 2)
        gd.addNumericField('UPPER_TH_CH%d' %chIndex, 254, 2)
    
    gd.addMessage("Segmentation channels (automated finding target region)",headerFont)
    gd.addStringField('Sum Channels for Segmentation', '%d; %d' %(1,nChannels) if nChannels>1 else '1', 0)
    
    
    gd.showDialog()
    if gd.wasCanceled():
        WaitForUserDialog("Analysis aborted by User").show()
        sys.exit("Analysis aborted by User")
    
    for chIndex in range (1,nChannels+1):
        inputCollection.addParameter('LOWER_TH_CH%d' %chIndex,gd.getNextNumber(),None,ParameterType.DOUBLE_PARAMETER)
        inputCollection.addParameter('UPPER_TH_CH%d' %chIndex,gd.getNextNumber(),None,ParameterType.DOUBLE_PARAMETER)
    
    segmentationChannelString=gd.getNextString()
    segmentationChannelIndexes=map(int,segmentationChannelString.split(';'))
    segmentationChannelIndexes=array(segmentationChannelIndexes,Integer)
    
    inputCollection.addArrayParameter("Segmentation Channels",segmentationChannelIndexes,None,ParameterType.INT_PARAMETER)
    
    return inputCollection


if __name__ == '__builtin__':


  
    print("#")

    print("# Tissue analysis")

    print("#")

    #

    # GET ANALYSIS PARAMETERS

    #

    
    analysisParameters=get_parameters()
    

    #

    # MAKE OUTPUT FOLDER

    #

    input_folder = analysisParameters.getParameterValue("Input Data Path")
    output_folder = input_folder.rstrip('\\/')+"--fiji"

    FileUtils.createCleanFolder(File(output_folder))
    

    # WRITE ANALYSIS PARAMETERS TO THE FILE
    ParameterXmlWriter().writeParametersToFile(analysisParameters,File(output_folder,"analysis_parameters.xml"))
    
    #

    # DETERMINE INPUT FILES

    #

    tbModel = createImageTableModel(_analysisPath = output_folder,_nChannels=analysisParameters.getParameterValue("Number of Channels"))
    tbModelRegions = createRegionTableModel(_analysisPath = output_folder,_nChannels=analysisParameters.getParameterValue("Number of Channels"))
    tbModel = determine_input_files(input_folder, tbModel,analysisParameters.getParameterValue("Input File Extension"))

    

    frame=ManualControlFrame(tbModel,False)

    frame.setVisible(True)

    

    #initialise Roi manager
    rm=RoiManager.getInstance()
    if rm is None:
        rm=RoiManager()
    
    #

    # ANALYZE

    #

    to_be_analyzed = analysisParameters.getParameterValue("Datasets to Analyse")
    if not to_be_analyzed=="all":

        close_all_image_windows()

        analyze(int(to_be_analyzed)-1, tbModel, tbModelRegions, analysisParameters)

        pass
    else:

        for i in range(tbModel.getRowCount()):

            close_all_image_windows()

            analyze(i, tbModel, tbModelRegions, analysisParameters)

        pass
    
    close_all_image_windows()
    
    tbModel.writeNewFile('analysis_summary_image.txt',True)
    tbModelRegions.writeNewFile('analysis_summary_regions.txt',True)
    
    print 'Analysis finished'
```
