## Supplementary material for "Multiomic and quantitative label-free microscopy-based analysis of *ex vivo* culture and TGFbeta1 stimulation of human precision-cut lung slices": ImageAnalysis: hPCLS-microscopy-image-analysisDescription.docx

Selective analysis of interstitial collagen in hPCLS:

Note:

- File format that can be analysed: Stitched tile acquired Z-stacks; all image file formats compatible with FIJI “bioformats” plugin
- For the script to work AutoMic_JavaTools-1.1.0-SNAPSHOT-19072016.jar should be placed/coppied in the plugin folder of FIJI

1) Open the Jython code in FIJI
2) Press “Run” to start the analysis
   a. Define the location of the images
      i. Note: as soon as the analysis starts a folder named as “....--fiji" is created. This folder contains all the processed data
3) Enter the number of channels that the z-stack have
4) Enter the number of ROIs that the stacks need to be divided into (e.g. 3X3 or more)
5) Enter the threshold values for signal in each channel (the values can be calculated before running the script separately)
6) A window with the defined number of ROIs will pop up, delete the ROIS that are not required (late on in the text file , the Boolean option will be 0 for this ROIs. Hence only, selected ROIs will be considered.
7) The image analysis generates two text files:
   a. “analysis_summary_image” with calculated values for different parameters for the whole z-tack (PCLS)
   b. “analysis_summary_regions” with calculated values for each ROI/ PCLS
8) Processed Binary (named as BW…) as well 8 bit images (named as GATED…) are generated for each channel of the z-stack, as part of the analysis as well
9) Second.Harmonic.Max.Proj folder contains processed image which is a sum of max projections of the two channels defined at the start of the analysis
10) The text file can be opened in excel and subjected to analysis as per the user’s needs

Different parameters and columns listed in the text file:

PBT.1_NUM: All the pixels present in the defined threshold

SumIntensity.1_NUM=Sum intensity of all the pixel in a channel

PBT.1AND2_NUM=Number of pixels common between respective channels

PBT.1OR2OR3_NUM= Number of pixels positive in either channels (multiple pixels positive in different channels only counted once)
